# Growth Mechanics: General principles of optimal cellular resource allocation in balanced growth

**DOI:** 10.1101/2022.10.27.514082

**Authors:** Hugo Dourado, Wolfram Liebermeister, Oliver Ebenhöh, Martin J. Lercher

**Affiliations:** Institute for Computer Science and Department of Biology, Heinrich-Heine Universität, 40221 Düsseldorf, Germany; Université Paris-Saclay, INRAE, MaIAGE, 78350 Jouy-en-Josas, France; Quantitative and Theoretical Biology, Heinrich-Heine Universität, 40221 Düsseldorf, Germany

**Keywords:** cell growth, self-replication, optimal resource allocation, growth economy, growth control

## Abstract

The physiology of biological cells evolved under physical and chemical constraints such as mass conservation, nonlinear reaction kinetics, and limits on cell density. For unicellular organisms, the fitness that governs this evolution is mainly determined by the balanced cellular growth rate. We previously introduced Growth Balance Analysis (GBA) as a general framework to model such nonlinear systems, and we presented analytical conditions for optimal balanced growth in the special case that the active reactions are known. Here, we develop Growth Mechanics (GM) as a more general, succinct, and powerful analytical description of the growth optimization of GBA models, which we formulate in terms of a minimal number of dimensionless variables. GM uses Karush-Kuhn-Tucker (KKT) conditions in a Lagrangian formalism. It identifies fundamental principles of optimal resource allocation in GBA models of any size and complexity, including the analytical conditions that determine the set of active reactions at optimal growth. We identify from first principles the economic values of biochemical reactions, expressed as marginal changes in cellular growth rate; these economic values can be related to the costs and benefits of proteome allocation into the reactions’ catalysts. Our formulation also generalizes the concepts of Metabolic Control Analysis to models of growing cells. GM unifies and extends previous approaches of cellular modeling and analysis, putting forward a program to analyze cellular growth through the stationarity conditions of a Lagrangian function. GM thereby provides a general theoretical toolbox for the study of fundamental mathematical properties of balanced cellular growth.

## Introduction

A core feature of microbial cells is self-replication – their ability to build a complete, identical cell exclusively out of the chemical compounds found in the environment. If a population of asynchronously replicating microbial cells grows exponentially in a constant environment, its self-replication can often be assumed to result from balanced growth, a non-equilibrium steady state in which every cellular component accumulates at the same rate in proportion to its total amount (1). For non-interacting microbes in a constant environment, the balanced growth rate is equivalent to fitness (2).

The cellular composition is thus often interpreted as an approximate solution to a problem of optimal allocation, driven by natural selection. Accordingly, theoretical methods estimating the optimal allocation are used as a reference to understand cellular composition in vivo (3–8).

At the whole-cell level, a mechanistic understanding of the quantitative principles that shape cellular balanced growth has been approached predominantly through methods collectively classified as constraint-based modeling (CBM). CBM approaches define a solution space of feasible cellular states (usually defined by reaction fluxes) based on simple, mechanistic constraints. The predominant constraint in CBMs is flux balance, encoded through a linear system of equations that constrain the space of allowed reactions fluxes **v** (9, 10),

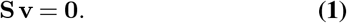

Here, **v** is a vector of reaction fluxes, i.e., reaction rates in units of [moles][time]^−1^[volume]^−1^. Each row of the stoichiometric matrix **S** corresponds to one metabolite, while each column corresponds to a metabolic reaction, with entries listing the corresponding stoichiometric coefficients of substrates (negative integers) and products (positive integers). Thermodynamics and physiological limits – such as a limited nutrient uptake capacity – are typically approximated through fixed upper and/or lower bounds on the modeled fluxes **v** (11). The most widely used CBM approach, flux balance analysis (FBA) (11, 12), obtains plausible physiological states by optimizing some objective function over the feasible flux vectors. Frequently, the objective function is the flux through a hypothetical biomass reaction *v*_*bio*_, which mimics the accumulation of precursors for macromolecules and the consumption of energy for their assembly during growth. Resource balance analysis (RBA) and metabolism and expression models (ME-models) go beyond FBA by aiming to model metabolism in its most general sense, with the ultimate goal of representing all chemical reactions that occur in a living organism (8, 13). In contrast to FBA, these methods take into account the burden of producing the macromolecules required for catalyzing each flux. They approximate the corresponding kinetic rate laws as linear relations between fluxes and the concentration of their catalysts, ignoring the dependence on reactant concentrations (except for dependencies on extracellular concentrations, which serve as model parameters).

All widely used CBMs (8, 11, 13, 14) are formulated as linear optimization problems, which can be solved efficiently even for genome-scale models with thousands of reactions. Accordingly, they are currently the most powerful tools to predict and understand realistic cellular models. However, by construction, these linear methods cannot capture the potentially complex nonlinear relationship between biochemical reaction fluxes – and hence cellular growth –– and the concentrations of reactants involved as substrates and products. Instead of accounting for nonlinear kinetics, these methods rely on linear, phenomenological assumptions.

Nonlinear CBMs (6, 15–19) account for constraints such as nonlinear kinetic rate laws, linking the concentration of metabolites to reaction fluxes. This link means that the metabolite concentrations are now an output of the model instead of an input. Molenaar et al. (6) introduced “self-replicator” models that maximize the cellular growth rate, with reaction fluxes that are limited by fundamental physiological constraints including mass conservation, nonlinear rate laws, and limited protein density. Importantly, these models are completely self-contained, in the sense that in order to grow and self-replicate, all of a model’s individual components have to be produced explicitly by the model itself. Instead of using a phenomenological “biomass reaction”, the constrained optimization of growth predicts the detailed cell composition, and all possible trade-offs in resource allocation can be accounted for from first principles.

Similar to RBA and ME models, self-replicator models include a “ribosome” reaction that produces the necessary proteins. The proteins can be classified into three categories: transport proteins in the cell surface, which exchange mass with the environment; enzymes, which catalyze internal metabolic reactions; and the ribosome itself, which catalyzes the internal protein production, and which for simplicity is assumed to be composed only of proteins. The study of models of this type relies on the numerical solution of nonlinear optimizations: while it in principle accommodates models with any number of reactions, actual presented models have small, highly simplified reaction networks (6, 15–19).

We have previously formalized a general framework for non-linear CBM and provided solutions for important special cases, an approach we termed growth balance analysis (GBA) (Fig. 1). GBA models are based on the self-replicator scheme. Instead of a fixed protein concentration, they consider a fixed combined mass density of their components. Optimal cellular resource allocation, as predicted by GBA models, emerges exclusively from quantitative biochemical and physical principles, including the intrinsic nonlinear nature of the underlying reaction kinetics. In general, the optimization of nonlinear models is a non-convex problem, frequently hampered by the existence of multiple local optima (20). Several studies have explored ad-hoc analytical solutions to convex, minimal nonlinear cell models consisting of up to three cellular reactions. Despite their simplicity, simulations with these schematic models are qualitatively consistent with the experimentally observed behaviour of actual cells (6, 15–19).

**Fig. 1.**
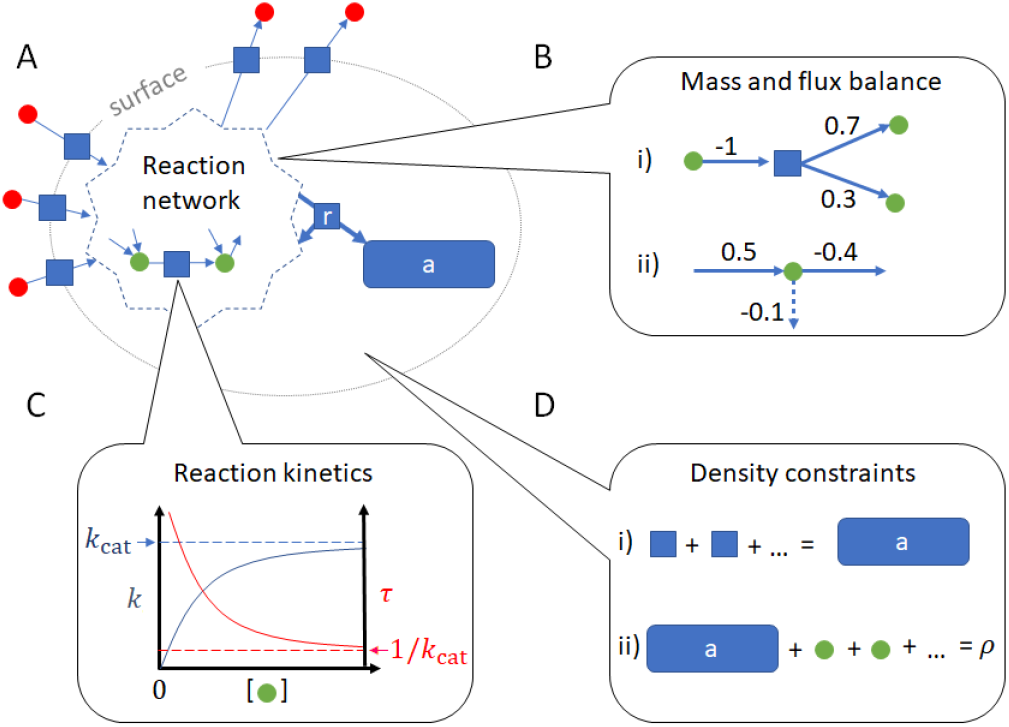
Constraints in a GBA model. **A)** In a GBA model, a cell exchanges external reactants (red circles) via transporters (blue squares at the cell surface); converts internal reactants (green circles) via enzymatic reactions (blue squares inside the metabolic network); and produces all proteins catalyzing the reactions (blue rectangle “a”) via a ribosome reaction “r”. The ribosome reaction consumes and returns metabolites to the metabolic network. In its strict sense, the metabolic network comprises the conversion of small molecules into energy and precursors for macromolecules. A model may also describe metabolism in its more general sense, including other enzymatic reactions such as those for DNA replication and transcription. **B)** All reactions in the model must conserve mass, a concept that comprises (i) mass balance within reactions: one unit of mass consumed (-1) results in one unit of mass produced (+1); and (ii) flux balance of reactant production and consumption, including the dilution by growth of all components (dashed arrow). **C)** Each reaction flux is catalyzed by a specific protein with turnover time *τ* (or equivalently, turnover rate *k* = 1*/τ*). *τ* is determined by kinetic rate laws and depends on the concentrations of reactants involved in the reaction; *k* = 1*/τ* has a maximal value *k*_cat_. **D)** Two basic density constraints govern the cellular interior: (i) the density of proteins “a”, and (ii) the total density *ρ*, which is the sum of all protein and metabolite concentrations.

Together with the GBA model framework, we introduced an analytical theory to study GBA models of any size in the special case that there are no redundant reactions, in the sense that the function of any reaction in the model cannot be copied by any combination of other reactions (i.e., the reaction matrix has full column rank). For this special case, we derived the necessary analytical conditions for optimal growth. Optimal states of arbitrarily complex models also have to satisfy these conditions, as they must always use non-redundant sets of active reactions (4, 21, 22). The necessary conditions provide a tool to understand important properties of cellular resource allocation at optimality, e.g., the protein allocation into ribosomes at different growth rates (4). On the other hand, the conditions cannot explain themselves why a particular set of non-redundant reactions is used at optimality. In the previously published GBA treatment, this choice has to be an input of the model. This is a major restriction, as real cells activate different sets of reactions depending on the growth conditions, and realistic mathematical models should be able to predict these sets.

Below, we analyze the growth optimization of arbitrary GBA models that may include redundant reactions, generalizing the results in Ref. (4). We provide the necessary analytical conditions for the optimality of each reaction flux, including the criteria for whether a reaction is active at optimality. We interpret these analytical conditions in terms of marginal costs and benefits of reactions with respect to their influence on growth, and quantify how changes in the model parameters and external conditions control the optimal growth rate.

## Results

We first present the notation and mathematical definition for growth optimization, including an objective function and constraints. We then reformulate the problem in terms of flux fractions as the only free variables, which greatly simplifies the subsequent analytical study. Finally, we explore the consequences that emerge from the necessary optimality conditions in terms of economics and control theory, and discuss their biological significance. Table 1 lists all symbols used below.

**Table 1.** Symbols used.

| Symbol | Description (units) |
| --- | --- |
| $A$ | growth adaptation coefficient |
| $\mathbf{b}$ | biomass fraction vector |
| $\mathbf{c}$ | reactant concentration vector ([mass][volume]<sup>-1</sup>) |
| $\mathbf{C}$ | control coefficient matrix |
| $\mathbf{f}$ | flux fraction vector |
| $\mathbf{E}$ | indirect sensitivity matrix ([time]) |
| $\mathbb{K}$ | set of kinetic parameters (various units) |
| $\mathcal{L}$ | Lagrangian ([time]<sup>-1</sup>) |
| $\mathbf{M}$ | mass fraction matrix |
| $\mathbf{p}$ | protein concentration vector ([mass][volume]<sup>-1</sup>) |
| $\mathbf{S}$ | stoichiometric matrix ([mol]) |
| $\mathbf{v}$ | flux vector ([mass][volume]<sup>-1</sup>[time]<sup>-1</sup>) |
| $\mathbf{x}$ | external concentration vector ([mass][volume]<sup>-1</sup>) |
| $\boldsymbol{\gamma}$ | vector with column sums of $\mathbf{M}$ |
| $\Gamma$ | growth control coefficient ([time]<sup>-1</sup>) |
| $\boldsymbol{\varepsilon}$ | direct sensitivity matrix ([time]) |
| $\boldsymbol{\phi}$ | proteome mass fraction vector |
| $\lambda_j$ | KKT multiplier for the protein constraint ([time]<sup>-2</sup>) |
| $\lambda_m$ | KKT multiplier for the metabolite constraint ([time]<sup>-1</sup>) |
| $\lambda_\rho$ | KKT multiplier for the density constraint ([time]<sup>-1</sup>) |
| $\mu$ | growth rate ([time]<sup>-1</sup>) |
| $\boldsymbol{\pi}$ | parameter vector (various units) |
| $\rho$ | mass density ([mass][volume]<sup>-1</sup>) |
| $\boldsymbol{\tau}$ | turnover time vector ([time]) |

| Index | Description |
| --- | --- |
| $a$ | all proteins |
| $e$ | enzymatic reactions |
| $i$ | internal reactants (including $m$ and $a$) |
| $j$ | reactions (including $s, e, r$) |
| $l$ | reactions (including $s, e, r$) |
| $m$ | metabolites |
| $r$ | ribosome reaction |
| $s$ | surface reactions (i.e., transport reactions) |

### Growth Modeling

We define a GBA model as the triple (**M, *τ***, *ρ*). The matrix **M** describes the mass fractions of internal reactants consumed and produced by each reaction; ***τ*** is a vector of catalytic turnover times for each reaction; and *ρ* is the combined mass concentration of all internal components. In the following paragraphs, we provide more detailed descriptions of the model constituents **M, *τ***, and *ρ*. Here and below, we use the term “reaction” to also encompass transport processes across the cell surface, which are “catalyzed” by transporter proteins or protein complexes.

**M** was first introduced in (4). It is constructed from the stoichiometric matrix **S** for the total, closed system, i.e., including rows for external reactants. We add a column “r” for the production of protein from precursors by the ribosome, as well as a row “a” corresponding to the total concentration of all proteins, following the procedure introduced by Molenaar et al. (6). We now first convert all entries to masses, by multiplying each row with the corresponding molecular mass. Because of mass balance, each column must then sum to 0. We next normalize each column such that the sum of negative entries equals −1 and the sum of positive entries equals +1. Now the entries correspond to the mass fractions of each reactant (rows) going into and out of each reaction (columns), as illustrated for the example in Fig. 1B. Finally, we reduce the normalized matrix to a matrix for an open system, by dropping all rows for reactants external to the modeled cell. For the remaining internal reactants, we will assume a quasi-steady state and thus enforce mass conservation.

As illustrated in Fig. 2, to simplify the notation for the following theoretical development, we partition the columns of **M** (indexed together by *j*) into index sets for reaction types: *s* for transport processes across the cell surface; *e* for internal enzymatic reactions; and r for the ribosome reaction, which is the only one that produces protein. We partition the rows of internal reactants (indexed together by *i*) into indices *m* for metabolites and a for total protein. We use the term “metabolites” in its more general sense, referring to any molecule in the cell that is not a protein. We distinguish vectors by using boldface, and vector components by using italics with the appropriate index, e.g., **c** is the column vector of all internal reactant concentrations, *c*^*i*^ are its components, and we use a lower index to indicate the components *c*_*i*_ of the row vector **c**⊤.

**Fig. 2.**
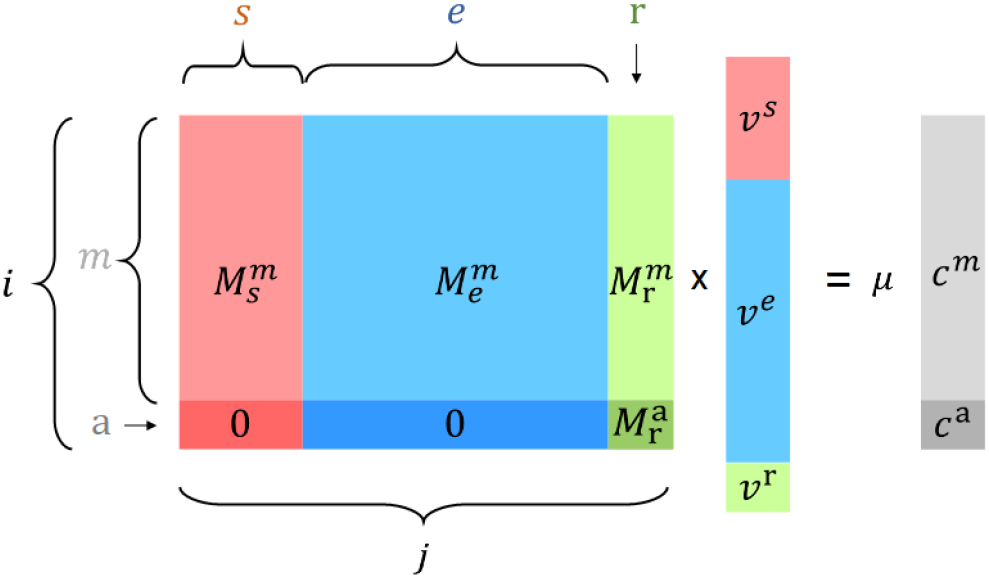
**Schematic overview of the mass conservation constraint Mv** = *μ***c**, determined by the mass fraction matrix **M**, the column vector of mass fluxes **v**, the growth rate *μ*, and the column vector of internal reactants mass concentrations **c**. The indices indicate partitions according to the type of reaction (columns of **M, v**) or reactant (rows of **M, c**). The index *i* = (*m*, a) correspond to rows for internal reactants, comprising rows *m* for metabolites, and a row “a” for the total mass concentration of all proteins. The indices *j* = (*s, e*, r) correspond, respectively, to transport proteins, enzymes, and the ribosome. We also use the index *l* for all reactions when necessary. Note the row “a” of **M** has only one nonzero entry 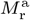, corresponding to the mass fraction of protein produced by the ribosome reaction r. Different colors indicate three different types of reactions: red (transporters), blue (enzymes), green (ribosome); and two types of reactants: light gray (metabolites), dark gray (total protein), resulting in six partitions of **M**.

The ribosome reaction represents the last step in protein synthesis, and is assumed to be catalyzed by a “ribosome” consisting entirely of protein. We here ignore the RNA components of the ribosome for simplicity, but it is possible to extend the modeled ribosome to a more realistic RNA-protein complex. In addition, the enzymatic reactions (*e*) could be extended so that they include details of protein translation that occur before the last, “ribosome” step (*r*) (5). Note that nonlinear genome-scale GBA models can be created from existing linear genome-scale models, by extending their stoichiometric matrix **S** with the addition of a ribosome reaction, normalizing it to **M** with the molecular masses, and adding the kinetic rate laws ***τ*** and density *ρ* (see below).

We assume that every reaction represented by a column *j* in **M** is catalyzed by a protein or protein complex with concentration *p*^*j*^ – specifically, a transporter (*s*), an enzyme (*e*), or the ribosome (r). The corresponding flux *v*^*j*^ is assumed to be proportional to *p*^*j*^, expressed as *v*^*j*^ = *p*^*j*^*/τ* ^*j*^(**c, x**). Thus, *τ* ^*j*^ is defined as the inverse of the usual metabolite-dependent factor in kinetic rate laws; SI text “Rate laws and kinetic parameters” provides a basic discussion of rate laws and the necessary kinetic parameters. **c** = (*c*^*m*^, *c*^a^)⊤ is the vector of internal reactant concentrations, comprising all metabolite concentrations *c*^*m*^ as well as *c*^a^, the combined mass concentration of all proteins. Hence, each *τ* ^*j*^ may depend not only on the concentrations of the substrates and products of the corresponding reaction, but also on inhibitors and regulatory metabolites not involved in the turnover itself. The transport processes *s* are the only reactions whose rate laws may depend on the external concentrations **x**.

Note that in accordance with the normalization of **M**, all concentrations of proteins *p*^*j*^ and reactants *c*^*i*^ throughout this work are in units of [mass *×* volume^−1^]. Fluxes ([mass *×* volume^−1^ *×* time^−1^]) and the kinetic parameters must then also be expressed in mass units, e.g., Michaelis constants *K*_m_ in [mass *×* volume^−1^] and turnover numbers *k*_cat_ in product mass per protein mass per time, resulting in [time^−1^].

*ρ*, the final constituent of GBA models, is the sum of all internal concentrations. We assume *ρ* to be constant, which is consistent with experimental data on *E. coli* across growth conditions and even across the cell cycle (23–25). The mass balances exploited for the normalization of **M** mean that all reactants involved in reactions must be accounted for in the model and hence be included in the value of *ρ*; e.g., in a realistic model water is a reactant in many reactions, so *ρ* corresponds in this case to the total cellular density (or buoyant density). Simplified models may instead include only dry mass components, so that both **M** and *ρ* consider only these. The equations below will be simplified by adopting the Einstein summation convention: a repeated lower and upper index denotes a summation over this index (often indicating a matrix multiplication), e.g., 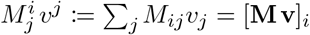. We also denote element-wise multiplication with repeated upper (or repeated lower) indices.

Mass conservation implies that in the mass fraction matrix **M**, each column sum 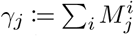 is zero if it involves only the consumption and production of internal reactants (indices *e*, r). In contrast, transport reactions (with indices *s*), which bring mass into and out of the modeled system, do not conserve mass, resulting in the equations

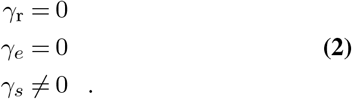

The property (2) guarantees mass conservation within reactions, an information that is not always fully encoded in the stoichiometric matrix **S** (see SI text “Mass balance and the stoichiometric matrix **S**”). While external reactants have no corresponding rows in **M**, their concentrations **x** may influence the turnover times *τ* ^*s*^ of transporters. We present examples of GBA models and R code for their numerical optimization in Dataset S1 (see SI text “Examples of GBA models and R code for numerical optimization”).

We are interested in the cellular physiology, defined through the concentration vectors **c, p** and the vector of reaction fluxes **v**, at balanced growth. For a given model specified by (**M, *τ***, *ρ*) and a given environment specified by **x**, balanced growth at the instantaneous rate *μ* is specified by the following constraints:

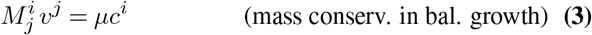

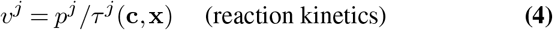

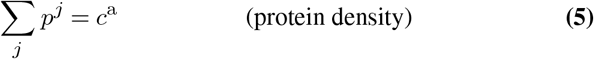

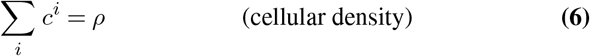

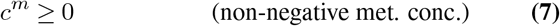

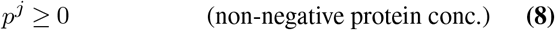

The model’s balanced growth property is captured by the right hand side of Eq. 3. We assume that the growth rate is always positive, *μ >* 0. Thus, for internal nodes with non-zero concentration (*c*^*i*^ ≠ 0), there is a necessary mass flow to offset the dilution through the associated volume growth at rate *μ*. Note that the total protein concentration *c*^a^ in Eq. 5 is not fixed, but is constrained by a row “a” in Eq. 3 that specifies mass conservation in balanced growth.

Below, we will be interested in optimal balanced growth, defined as growth at the maximal possible rate *μ* given the above constraints. From Eq. 3-5, we see that the variables (**c, p, v**) are highly interdependent. The above formulation does not lend itself to expressing *μ* as an explicit function of these variables, which makes it not ideal for numerical or analytical studies. If one can find a mathematically equivalent formulation based on fewer, independent variables, then this would facilitate the use of the KKT conditions, analogous to how generalized coordinates facilitate the solution of problems in Lagrangian mechanics (26). Thus, we next focus on a corresponding reformulation of the optimization problem. This formulation will apply to all balanced growth states, and only later we will use it to examine optimal balanced growth states.

### A Reformulation in Terms of Flux Fractions f

Our guiding thought below is that there can be a correspondence between cell states at different growth rates, which can be expressed in the form of scaling relations. These scaling relations extend the mass fraction scaling of **M** to fluxes and concentrations. Specifically, we define *biomass fractions*

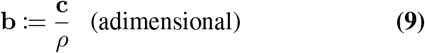

(equivalent to **c** = *ρ* **b**, since *ρ >* 0), which express concentrations as fractions of the total cellular density; and we define *flux fractions*

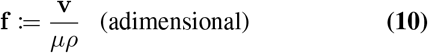

(equivalent to **v** = *μρ* **f**, since we assume *μρ >* 0), which express fluxes as fractions of the net mass uptake – i.e., the net growth – of the cell, *μρ*. Thus, each flux fraction *f* ^*j*^ describes the activity of reaction *j* relative to the total cellular mass production. Importantly, Eq. 3 describing mass conservation in balanced growth does not depend explicitly on *μ* anymore when written in terms of **f** and **b**:

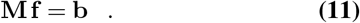

This equation also implies that the mass fractions **b** are uniquely determined by the flux fractions **f**, independently of *μ*. Conveniently, this unique dependence also means that we can express the turnover times as functions of only **f** and the fixed parameters *ρ*, **M**, and **x**:

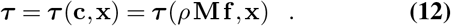

In the following discussion, we mostly focus on the dependence of ***τ*** on **f**, and for simplicity of notation we do not state the dependence of ***τ*** on the fixed parameters (*ρ*, **M, x**) explicitly. Importantly, ***τ*** does not depend explicitly on *μ*, which otherwise would cause a recursion problem when further expressing the growth rate *μ* in terms of only **f** and ***τ*** (**f**), as we will see below. The resulting dimensionality reduction of the solution space not only simplifies the analytical considerations below, but also potentially facilitate numerical optimizations (27).

From Eqs. 4 and 10, we obtain the expression for protein concentrations in terms of **f**, *μ*, and *ρ*,

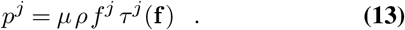

The combined mass fraction of all proteins in the cell, *b*^a^, is the sum of all *p*^*j*^ in the last equation, divided by *ρ*:

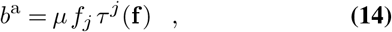

where we indicate summation by combining the lower index in *f*_*j*_ with the upper index in *τ* ^*j*^. Thus, we can calculate the total protein mass fraction during balanced growth from *μ* and **f**, based only on reaction kinetics. However, through Eq. 11, the same total protein mass fraction is also related to **f** through the corresponding row “a” in **M**:

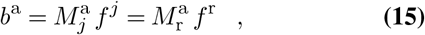

where the second equality reflects our assumption that the “ribosome” reaction r is the only one producing proteins, so that 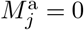 for *j* ≠ r. Equating the right hand sides of the previous two equations and solving for *μ* (with *b*^a^ ≠ 0 ⇒ *f*_*j*_ *τ*^*j*^(**f**) ≠ 0), we get the *growth function*

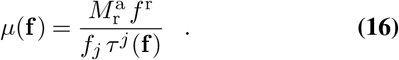

Thus, the growth rate becomes an explicit function of only the flux fractions **f**. *μ* still depends on the fixed parameters *ρ*, **M**, and **x** through the functions ***τ*** = ***τ*** (*ρ***Mf**, **x**). Note that if fluxes **v** were used instead of the flux fractions **f**, then ***τ*** (**c, x**) = ***τ*** (**Mv***/μ*, **x**), which would cause a recursion issue when defining the growth rate as a function of **v** and ***τ*** following the same procedure. In that case, one is forced to account for **c** as separate variables, thereby increasing the dimensionality of the problem. The same recursion issue occurs when formulating the problem in terms of protein concentrations.

From now on, we will consider **b** (Eq. 11) and ***τ*** (Eq. 12) as functions of **f**, and treat **f** as the only free variables. After writing the growth rate *μ* as a function of **f**, we now do the same thing for our remaining constraints, so now we have much fewer variables and constraints.

In the scaled variables, the density constraint (Eq. 6) is reduced to

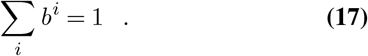

Using the balanced growth Eq. 11, we can rewrite this constraint in terms of flux fractions **f**. We see that in balanced growth, the density constraint (Eq. 17) is equivalent to a flux balance on the cell surface,

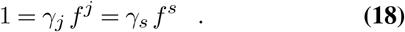

The second equality comes from Eq. 2: only the columns *s* sum up to non-zero values *γ*_*s*_, so only transport fluxes *f* ^*s*^ are limited by this constraint. The nature of this constraint as a global mass balance becomes more evident if we multiply the whole expression by *μρ*: the net mass uptake *γ*_*s*_ *v*^*s*^ going through the cell surface must equal the rate of biomass production *μρ*.

Any solution to the growth function (Eq. 16) automatically respects internal mass conservation, protein density and the kinetic constraints: for any given vector **f**, *μ*(**f**) returns the unique growth rate satisfying these constraints (which also depend on *ρ* through ***τ*** = ***τ*** (*ρ* **Mf**)). The flux balance at the cell surface is enforced separately by Eq. 18 on transporters, making these fundamentally different from enzymatic and ribosome reactions. In particular, for a model with only one transporter *s*, Eq. 18 already determines the scaled uptake rate *f*_*s*_ = (*γ*_*s*_)^−1^. With two transport fluxes, one flux is uniquely determined by the other; a simple example would be a model that only has transporters for glucose uptake and CO_2_ excretion (see example model “C” in SI text “Examples of GBA models and R code for numerical optimization”). More generally, Eq. 18 can be used to uniquely determine one transport flux fraction in terms of the others, reducing the number of variables by one. For clarity of presentation, however, we will keep Eq. 18 as a separate constraint and not eliminate any variable, until the introduction of growth control coefficients in the corresponding section.

Finally, writing the non-negativity constraints on proteins and metabolite concentrations in terms of **f** results in

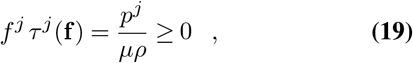

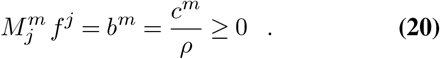

The non-negativity of the total protein concentration does not have to be imposed explicitly, as it already follows from the non-negativity of individual proteins. We are now in the position to provide a concise formulation of growth rate optimization in terms of flux fractions **f**. Combining Eqs. 16, 18, 19, 20, the optimal growth problem for a given environment **x** becomes

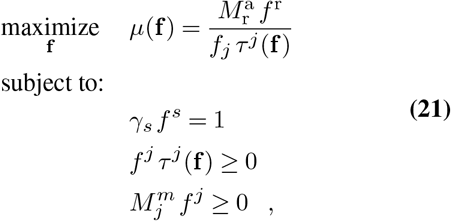

where the turnover times *τ* ^*j*^ are functions that depend on **f** and on the parameters *ρ*, **M, x**, *τ* ^*j*^ = *τ* ^*j*^ (*ρ* **Mf, x**). If the rate laws have a general functional form, these functions will be parameterised by the set kinetic parameters 𝕂. After solving this optimization problem, all original cellular variables (unscaled fluxes as well as unscaled metabolite and protein concentrations) can be easily reconstructed from **f**. In the following, we will refer to ***π*** as the vector of parameters that define the optimization problem, which includes **M**, *ρ*, and **x**, as well as the elements of 𝕂. The parameters in ***π*** are considered fixed until the section “Growth Control and Adaptation”, where we study the sensitivity of optimal growth to marginal changes in the components of ***π***.

### Growth Analysis

Next, we utilize the problem reformulation to derive general necessary conditions of cellular systems at optimal balanced growth, independent of model specifics. First, for each reaction, we will derive explicit expressions for shadow prices in the optimal state. We then derive equations for the state variables **f** themselves, which must hold in any optimal state. This development constitutes a generalization of our previous analytical approach to GBA (4), which was restricted to models with matrices **M** of full column rank. The latter condition is not generally satisfied by realistic cellular models, as many cellular biochemical reactions are structurally redundant, i.e., their columns in **M** are linearly dependent on other columns. Optimal growth states always have non-redundant active reactions (i.e., the *active* **M** has full column rank) (4, 21, 22), but this optimal set of active reactions is generally not known *a priori*. In contrast, the following analysis in terms of flux fractions is valid for any **M** of arbitrary size and rank.

For the following, we emphasize that the state of our system is completely determined by scaled fluxes *f* ^*j*^, which serve as independent variables. All other variables are fully dependent on them: the unscaled fluxes **v**, the scaled and unscaled concentrations **b, c**, and **p**, the reaction times ***τ***, and the growth rate *μ*.

All following analyses benefit from knowing the system’s sensitivity to small changes of each of the independent variables *f* ^*j*^. The partial derivatives of the system’s properties **c**(**f**), **v**(**f**), **b**(**f**), ***τ*** (**f**), **p**(**f**), and *μ*(**f**) with respect to each *f* ^*j*^ provide explicit expressions for sensitivity coefficients similar to the ones introduced in Metabolic Value Theory (28, 29), based on the original concepts of Metabolic Control Analysis (MCA) (9, 10). A unique feature of the present treatment arises from the system of equations in Eq. 11, which determines the linear dependence of biomass fractions **b** on **f**, so that the partial derivative of *b*^*i*^ with respect to *f* ^*j*^ is given simply as

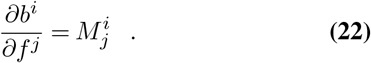

Via the chain rule of differentiation, this expression also determines the partial derivatives with respect to *f* ^*j*^ for any functions of *b*^*i*^. A case of particular interest in the following discussions is the vector of turnover times ***τ*** = ***τ*** (**c, x**) = ***τ*** (*ρ* **b, x**) = ***τ*** (*ρ* **Mf, x**). We first define the (direct) *time elasticities* (*elasticities* in short), the sensitivity of each turnover time *τ* ^*l*^ (**c, x**) = *τ* ^*l*^ (*ρ* **b, x**) with respect to each biomass fraction *b*^*i*^, as

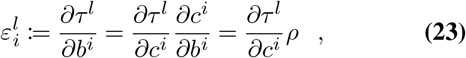

where we used the chain rule of differentiation in the first equality and Eq. 9 in the second. We then use the direct elasticities *ε*^*l*^ to express the sensitivity of 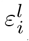 to a change in a flux fraction *f* ^*j*^, defined as the *indirect time elasticity* matrix **E** (or *indirect elasticity* in short), with entries

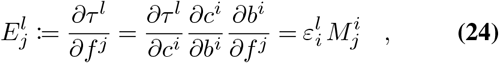

where we sum over *i* and used Eq. 22 in the last equality. In the following discussion, we assume that the kinetic rate laws do not depend on the total protein concentration *c*^a^, meaning 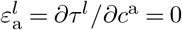 for all reactions *l*. That would be different if, for example, one accounts for the macromolecular crowding effects via kinetic rate laws (30). The indirect elasticities **E** and direct elasticities ***ε*** share some resemblance with the Jacobian and elasticity matrices defined in Metabolic Value Theory and MCA, although we do not intend to explore the exact relationships in this work. For an example of direct and indirect elasticities, where ***τ*** follows a simple Michaelis-Menten rate law, see SI text “Rate laws and kinetic parameters”.

In the remainder of this paper, we will explore three complementary types of analyses of GBA systems. First, in the growth optimality section we will state the analytical conditions necessary for an optimal state **f** *. Second, in the growth economy section we will calculate the sensitivity of a (not necessarily optimal) growth rate *μ* to small changes in each **f**, which we interpret in economic terms as marginal values of reactions. Third, in the growth control and adaptation section we will estimate the sensitivity of the optimal growth rate *μ*^*^ to small changes in the previously fixed parameters ***π***. In each of these analyses, the sensitivity measures captured by **E** will appear naturally in the results.

### Growth Optimality

We next calculate the necessary analytical conditions for the optimal growth state. This calculation extends our previous analytical approach, which was restricted to GBA models with matrices **M** of full column rank (4), to general GBA models with arbitrary matrices **M**, facilitated by the reformulation of the GBA problem in terms of flux fractions **f**. We approach this problem by studying the Karush-Kuhn-Tucker (KKT) conditions (31, 32), which generalize the method of Lagrange multipliers by also accounting for inequality constraints, here present due to the non-negativity of concentrations. To simplify the presentation in this section, we here account explicitly only for the non-negativity of protein concentrations, but not for the non-negativity of metabolite concentrations. Under the reasonable assumption that metabolites with zero concentration do not participate in any active reactions, the resulting necessary conditions are also necessary when accounting for this latter constraint; the full calculations can be found in the Materials and Methods section.

We define the Lagrangian ℒ (**f**, ***λ***) for a given GBA model (**M, *τ***, *ρ*) and external concentrations **x** as

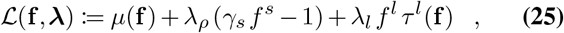

where we sum over *l* in the last term. The *KKT multipliers* ***λ*** are auxiliary variables used to find the optimal state, but also encode important economic and control information about the system at optimality, as we will see later. *λ*_*ρ*_ relates to the density constraint connected to **f** via the flux balance at the surface (Eq. 18); the *λ*_*l*_ relate to the non-negativity of proteins (Eq. 19). The necessary KKT conditions include the primal feasibility conditions given by these two equations and

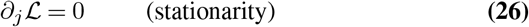

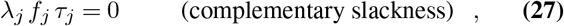

where ∂_*j*_ := ∂*/*∂*f* ^*j*^ indicates the partial derivative with respect to *f* ^*j*^.

The stationarity conditions can be solved for the corresponding optimal multipliers *λ*_*j*_, resulting in

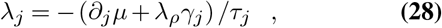

where *λ*_*ρ*_ is the optimal value for the density multiplier. After an element-wise multiplication of both sides of Eq. 28 with *f*_*j*_*τ*_*j*_, we can use the complementary slackness (*λ*_*j*_*τ*_*j*_*f*_*j*_ = 0) to get

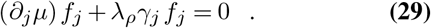

Now summing the last equation over all *j* and using the primal feasibility (Eq. 18) results in

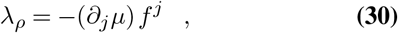

where we have a summation over *j*.

From Eq. 16, the partial derivative of *μ* with respect to *f* ^*j*^ is

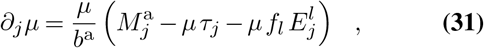

where we used the identity 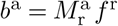 from Eq. 15. Substituting these derivatives into Eq. 30) results in an expression for the optimal value of the density constraint multiplier *λ*_*ρ*_ in terms of **f** only,

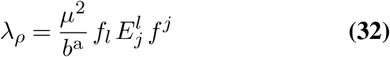

(see Materials and Methods for the detailed calculations). Note that in the last equation, *j* and *l* do not refer to specific reactions but instead indicate summations. When we further consider that only transporters have a nonzero column sum (Eq. 2), we get an equivalent expression that highlights the particular dependence of *λ*_*ρ*_ on the reactions directly connected to the transport reactions (see SI text “The dependence of *λ*_*ρ*_ on transporters”).

Combining Eq. 28, 31, 32, we can now express each multiplier *λ*_*j*_ explicitly in terms of the flux fractions **f** at optimality, resulting in slightly different expressions for ribosomal, enzymatic, and transport reactions:

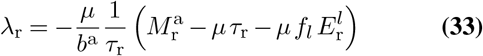

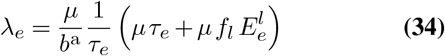

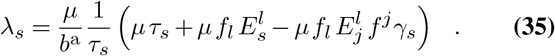

By inserting these expressions into the complementary slackness conditions (Eq. 27), we can now solve for **f**, which results in the three *balance equations* for ribosomal, enzymatic, and transport reactions:

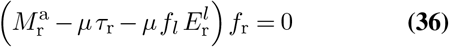

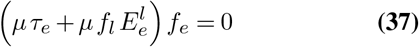

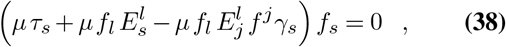

where we simplified the expressions by exploiting that *μ, b*^a^, *τ*_*j*_ ≠ 0. The balance equations are necessary equality conditions for the optimal growth state **f** * of the optimization in Eq. 21, and generalize Eq. 10 in (4). Note that in all cases, a reaction with nonzero flux *f* ^*j*^ requires that the corresponding term in parentheses (i.e., the corresponding *λ*_*j*_) is equal to zero. Conversely, if the term in parentheses is different from zero (*λ*_*j*_ ≠ 0), then the reaction cannot carry flux at optimality (*f* ^*j*^ = 0). In particular, the ribosome evidently needs to be active for balanced cellular growth, so *λ*_r_ = 0 must always hold in optimal states. The KKT multipliers *λ*_*j*_ can be understood as *shadow prices* of the scaled protein concentrations *τ* ^*j*^*f* ^*j*^ = *p*^*j*^*/*(*μρ*) (protein concentration per biomass production *μρ*). These shadow prices must be zero for active reactions at optimality, as otherwise growth rate could be increased by changing the corresponding protein concentration.

We may also express the balance equation for each reaction *j* in the usual, unscaled variables **p** (protein concentrations), **v** (fluxes), and **c** (reactant concentrations, including metabolites and total protein), by using Eqs. 4,9,10,11 (see Materials and Methods)

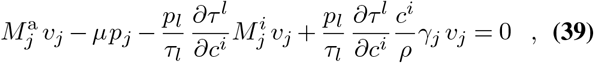

where we sum over indices *l* and *i*. We now continue our analysis in terms of the flux fractions **f**, since these are the variables of the optimization problem (Eq. 21). However, we keep in mind that the same change of variables to **p, v, c** is possible in all the following equations, as done for Eq. 39.

### Growth Economy

As growth rate is closely related to fitness (2), it makes sense to view growth rate as the primary value of the cellular economy. In this subsection, we will thus explore the economy of balanced cellular growth, by asking how a small change in the state variables *f* ^*j*^ affects the growth rate *μ* of any optimal or non-optimal state. Below, we will see that the necessary conditions of optimal growth specify that the marginal costs and benefits of each flux must be perfectly balanced.

We define the *marginal value* of flux *j* as the partial derivative ∂_*j*_*μ*, which quantifies the marginal gain in growth rate resulting from a small increase in *f* ^*j*^. From Eq. 31, we see that the marginal value can be expressed naturally as a multiple of the growth rate per mass fraction of protein in biomass, *μ/b*^a^. As we will see next, the corresponding adimensional factor –– the term in parentheses in Eq. 31 – quantifies different types of costs (when negative) and benefits (when positive) of reaction *j* in terms of its influence on protein allocation,

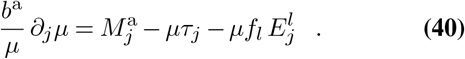

The first summand quantifies how a marginal increase in *f* ^*j*^ increases the total protein fraction in the cell density *b*^a^ = *c*^a^*/ρ* (see Eq. 5),

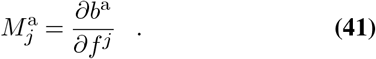

We name this contribution to the normalized marginal value the *marginal protein production*. As we assume that the ribosome reaction is the only reaction that consumes or produces protein, this reaction (*j* = r) is the only one with a nonzero (and positive) marginal production benefit.

To interpret the remaining summands, we remember that an individual protein’s mass fraction in the cellular density can be expressed as *p*^*l*^*/ρ* = *τ* ^*l*^*v*^*l*^*/ρ* = *μτ* ^*l*^*f* ^*l*^. The last two terms in Eq. 40 quantify the combined decrease of individual protein fractions in cellular density (*p*^*l*^*/ρ*) caused by a marginal increase in *f* ^*j*^ at fixed *μ*,

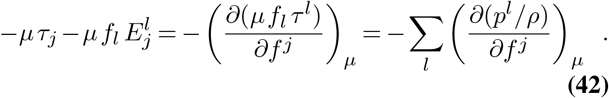

Here, the first summand quantifies the change in *p*^*l*^*/ρ* at fixed turnover times, which is evidently non-zero only for the enzyme catalyzing the perturbed flux *j* itself. We name this term, −*μτ*_*j*_, the *marginal (protein) investment* into *j*. The final summand quantifies the local change of the individual protein concentrations that must occur to compensate the changes in the turnover times (quantified by the indirect elasticity **E**), themselves caused by changes in metabolite concentrations forced due to flux balance. We name it the *marginal (protein) opportunity* of *j*, as it is related to opportunity costs and benefits in economics. For the typical case of reactions running in the forward direction (*f* ^*j*^ *>* 0), *τ* ^*j*^ is positive, and thus the marginal investment into *j* is negative, representing a cost. If all fluxes are non-negative, beneficial decreases in turnover times correspond to negative *E*, resulting in positive marginal opportunity (i.e., marginal opportunity benefit).

We can now summarize our insights about cellular economy, in particular about changes in the growth rate *μ* in response to changes in a flux *f* ^*j*^. The first and second terms in Eq. 40 are simple, direct consequences of the flux change: the marginal production benefit, an increase in protein production if *f* ^*j*^ is the ribosome flux; and the marginal investment, an increase in the protein concentration required to sustain an increased *f* ^*j*^. The third term in Eq. 40, the marginal opportunity, is more interesting, though equally easy to understand. As a simple consequence of mass conservation (Eq. 11), a change in *f* ^*j*^ while keeping all other fluxes fixed must result in changes in the concentrations of all reactants consumed or produced in the corresponding reaction. These concentration changes modify the turnover times *τ*_*l*_(**c**) of all reactions *l* whose kinetics depend on them, either because they are directly connected to those reactants or because they act as inhibitors or activators; see Fig. 3 for an example. Keeping the corresponding fluxes *f* ^*l*^ constant requires matching changes in the concentrations *p*_*l*_ of the catalyzing proteins (Eq. 4). This total amount of “protein saved” due to a change in *f*_*j*_ is quantified by − 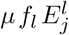.

**Fig. 3.**
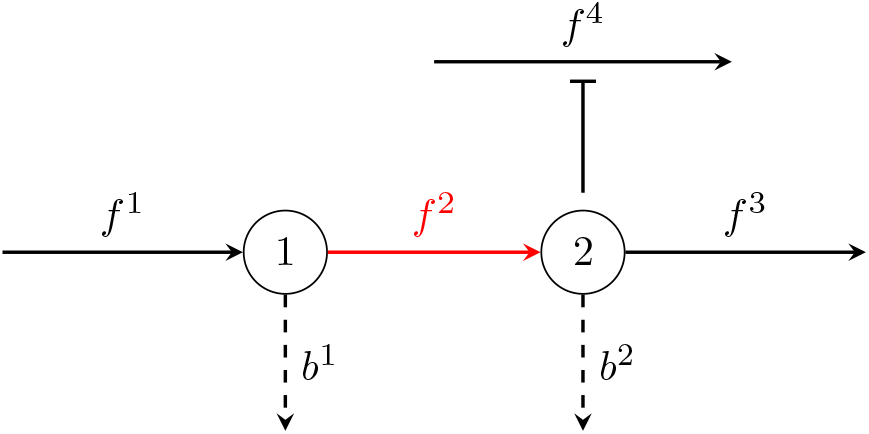
The dependence of marginal opportunity on the reaction neighborhood. The figure shows a simple example of a reaction (*j* = 2, red) that is directly connected to two metabolites (*m* = 1, 2) and thereby to two other reactions *j* = 1, 3. Reaction *j* = 2 is also connected indirectly to reaction *j* = 4 by inhibiting it through metabolite *m* = 2 (indicated by the blunt arrow ⊺). The marginal opportunity of *j* = 2 is − 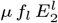, where 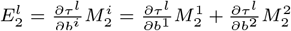 is determined by a marginal change in *f* ^2^ while keeping all other *f*^*l*^ fixed, causing (i) an inevitable change in *b*^1^, *b*^2^ due to the flux balance (Eq. 11); which by consequence causes (ii) an inevitable change in *τ* ^1^, *τ* ^2^, *τ* ^3^, *τ* ^4^, as these are functions of *b*^1^, *b*^2^; which finally causes (iii) an inevitable change in *p*^1^, *p*^2^, *p*^3^, *p*^4^ due to the kinetic constraints (Eq. 4) at fixed *v*^1^, *v*^2^, *v*^3^, *v*^4^ (determined by the fixed flux fractions and growth rate (Eq. 10)). The example also shows how the information about mass conservation and reaction kinetics is completely built into the definition of the growth function (Eq. 16).

The above results confirm the often postulated central role of proteins in the cellular economy (33–35). While the measure of cellular economic value may be the growth rate itself, protein concentrations constitute the general currency in which we can express the contributions of cellular subsystems. We can highlight this central economic role of proteins further by relating the marginal values ∂_*j*_*μ* – changes in growth rate in response to flux changes – to changes in the allocation of proteome fractions *ϕ*^*l*^ := *p*^*l*^*/c*^a^ :

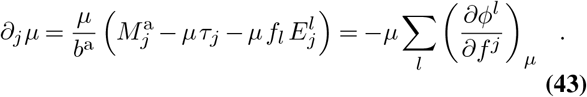

The second equality follows directly from taking the derivative of 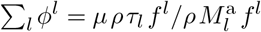 with respect to *f* ^*j*^ at constant *μ*.

We can now look at the balance equations at optimal growth from an economic perspective. For the ribosome and active enzymatic reactions, a zero marginal value ∂_*j*_*μ* = 0 also means a zero shadow price, *λ*_*j*_ = 0 (Eqs. 33, 34) – so the reaction is optimal, and growth cannot be accelerated by increasing or decreasing *f* ^*j*^ by a small amount. This insight provides an intuitive interpretation for the balance equations for the ribosome (Eq. 36) and for all active, internal enzymatic reactions (Eq. 37). An exception are only the transporters. In contrast to all other flux fractions, their shadow price (Eq. 35) depends both on their marginal value and on their *marginal biomass production*, 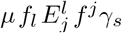 (a cost when negative and a benefit when positive).

For active enzymes with zero marginal value – and thus for all active enzymes at optimality (Eq. 37) – Eq. 31 simplifies to

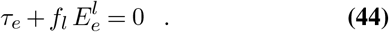

This simple relationship shows that at optimality, the marginal investment into *e* should perfectly balance its marginal opportunity. As the last equation involves only the neighborhood of *e* (defined as all reactions *l* such that 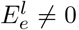), we can study such relationships at optimality locally, without full knowledge about the entire reaction network. We thus do not need the entire matrix **M** or complete knowledge of parameterized turnover time functions in the vector ***τ***.

In the preceding subsection, we studied how any optimal or non-optimal growth rate *μ* is sensitive to marginal changes in one of the flux fractions, resulting in an economic understanding of marginal flux values in terms of their relationship with protein allocation. We next reinterpret some of these results from the perspective of control theory, and turn to a complementary problem that focuses on the sensitivity of the optimal growth rate to changes in the model parameters and external concentrations.

### Growth Control and Adaptation

We are first interested in the total control that each *f* ^*j*^ has on the (optimal or non-optimal) growth rate *μ*, accounting also for the density constraint limitation. In order to do that, we choose one active transport reaction *s*^′^ and express its corresponding *f* ^*s*′^ ≠ 0 as a function of the other fluxes via the density constraint (Eq. 18),

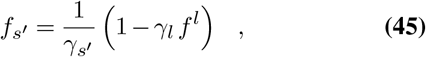

where *l* ≠ *s*^*′*^ sums over all other reactions. Thus,

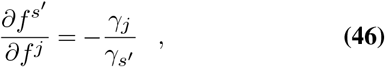

which is non-zero only if *j* is also a transport reaction (so *γ*_*j*_ ≠ 0). We now define the *Growth Control Coefficients* Γ_*j*_ as

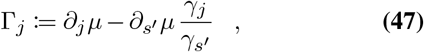

where the first term quantifies the growth change caused by *f* ^*j*^ itself, and the second term quantifies the growth change caused by a change in *f* ^*s*′^, itself changed due to the changed *f* ^*j*^ and the density constraint. Note that for the ribosome and enzymatic reactions, their growth control coefficient is simply their marginal value, since *γ*_r_ = *γ*_*e*_ = 0. For models with only one transport reaction *s*, Γ_*s*_ = 0, since *f*_*s*_ = (*γ*_*s*_)−1 is fixed by the density constraint and cannot be changed. Conveniently, this is also captured by Eq. 47. If *s*^*′*^ is optimal, *λ*_*s*′_ = 0, and Eq. 28 determines ∂_*s*_′ *μ* = −*λ*_*ρ*_*γ*_*s*′_, so in that case

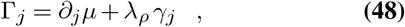

and Eqs. 36, 37, 38 are thus equivalent to

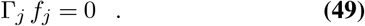

We may also see the optimal condition for enzymes (Eq. 44) in terms of protein concentrations, by multiplying it element-wise with *v*_*e*_ (so it is also valid now for inactive enzymes),

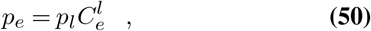

where we defined

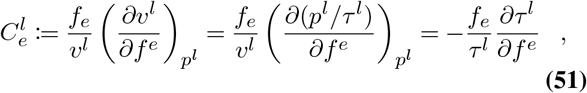

via Eq. 4 and using partial derivatives at fixed *p*^*l*^. 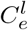 can been seen as (scaled) *control coefficients* (CC), analogous to (scaled) control coefficients in MCA (9, 10). This result is analogous to how enzyme concentrations and their respective CC relate at optimal fluxes constrained by a fixed total enzyme concentration (36) (see SI text “Optimal enzyme concentrations and control coefficients” for a detailed discussion). For an example of control coefficients where ***τ*** follows a simple Michaelis-Menten rate law, see SI text “Rate laws and kinetic parameters”.

We now explore the sensitivity of the optimal growth rate to changes in one parameter *π* in the vector ***π***. The growth problem (Eq. 21) is constrained by the parameters ***π***, including the arguments necessary to determine the turnover times ***τ*** at given **f**. This means that any marginal change in one of the parameters ***π*** would lead to changes in the solution **f** * of the optimization (Eq. 21). In this sense, the parameters ***π*** can be understood as control variables, while the corresponding optimal state **f** *, and its functions *μ*^*^ = *μ* (**f** *), **v**\* = **v**(**f** *), **p**\* = **p**(**f** *), and **c**\* = **c**(**f** *) are the response variables. Figure 4 summarizes these relationships.

**Fig. 4.**
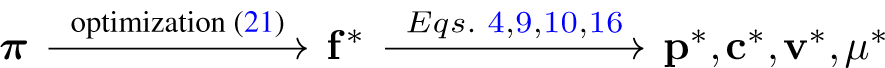
The parameters ***π*** and their control on the optimal cellular state **f**^*^.

Because growth rate is closely related to fitness, we are also particularly interested in how marginal changes in one of the previously fixed parameters ***π*** affect the *optimal* growth rate *μ*^*^ (37). We can estimate this effect directly via the envelope theorem (4, 38), by effectively considering the *optimal* state **f** * as fixed and treating the parameters ***π*** as the new independent variables, making it unnecessary to calculate the new optimal state after the parameter change. To do that, we first simplify the problem by assuming that these marginal changes have no effect on which reactions are active, so we simplify the Lagrangian (Eq. 25) by ignoring the inequality constraints; note that in this case only the objective function *μ* can be influenced by parameter changes, since the density constraint only depends on **M**, whose entries cannot be changed continuously. Second, we can think about the optimal growth rate *μ*^*^ as a function of the parameters *μ*^*^(***π***) := ℒ (**f** *(***π***), ***λ***^*^(***π***), ***π***), so the the total change d*μ*^*^*/*d*π* induced by a marginal change in a parameter *π* can be calculated via the chain rule

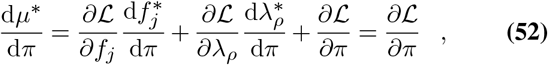

where the last equality comes from ∂ℒ */*∂*f* ^*j*^ = ∂ℒ */*∂*λ*_*ρ*_ = 0 according to the stationarity (Eq. 26) and primal feasibility (Eq. 18) at an optimal state.

We now define *growth adaptation coefficients A* as the relative change in the *optimal* growth rate *μ*^*^ in response to a small, relative change in one control variable *π*

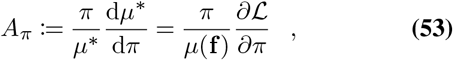

where here and in the rest of this section **f** is to be understood as the *optimal* state before the change in the parameter *π*. Note that in the following discussion, the parameters *π* of interest only influence ℒ via the objective function *μ*, so the partial derivatives ∂ℒ */*∂*π* are simply evaluated as ∂*μ/*∂*π* at fixed **f**.

For direct changes in the turnover times *τ* ^*j*^ (e.g., through changing the corresponding 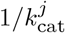), the growth adaptation coefficient is calculated by evaluating the growth function *μ* and its partial derivative at fixed **f**,

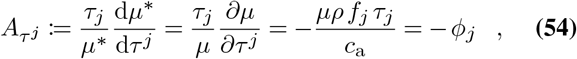

where we effectively treated *τ* ^*j*^ as a variable in the growth Eq. 16, and *ϕ*_*j*_ = *p*_*j*_*/c*_a_ is the optimal proteome fraction allocated to reaction *j* before the change in *τ* ^*j*^. This result is consistent with the observation that drugs targeting the most highly expressed catalysts, such as the ribosome, have the strongest effects on cellular growth rates (5, 39).

For changes in some external parameter such as a concentration *x*, the growth adaptation coefficient is again calculated by evaluating the growth function *μ* and its partial derivative at fixed **f**, and using the chain rule of differentiation we obtain

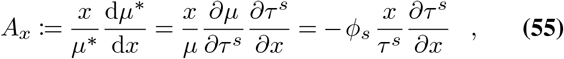

where we have a summation over *s* (only transporters *s* have kinetic rate laws depending on external concentrations). According to Eq. 55, the growth adaptation coefficient of an external concentration *x* is simply the sum over the “scaled elasticities” 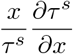 of the transporters of *x*, weighted by the optimal proteome fractions *ϕ*_*s*_ allocated to each *s* before the change in *x*. This result gives an explicit quantitative estimation on which external concentrations should be changed in order to cause the most change in the optimal growth rate. This equation may hence provide a useful tool for improving the growth media environment for industrial cell cultures, and for quantifying the effect of drugs aimed at decreasing the growth of pathogens and cancer cells. If the turnover times ***τ*** depend explicitly on other external parameters, such as pH and temperature, growth adaptation coefficients can be calculated and interpreted exactly as in Eq. 55.

The growth adaptation coefficient with respect to the mass density *ρ*, assuming it affects turnover times ***τ*** only through reactant concentrations **c**, reads

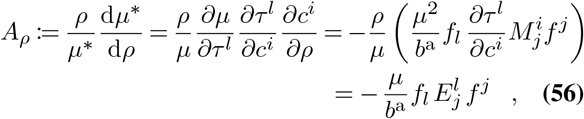

where 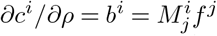 according to Eqs. 9,11, and the last equality comes from the definition of the indirect elasticity (Eq. 24). From this expression and *λ*_*ρ*_ in Eq. 32, we see that −*λ*_*ρ*_ = *μA*_*ρ*_; at optimality, the negative Lagrange multiplier for the density constraint, −*λ*_*ρ*_, quantifies the absolute increase in growth rate caused by a marginal increase in *ρ*, given by *μ* itself times the proportional change, *A*_*ρ*_. Thus, the extra term in the shadow price of transporters (compare *λ*_*s*_ in Eq. 35 to *λ*_*r*_ and *λ*_*e*_) quantifies the growth rate benefit gained by allowing the violation of the density constraint (Eq. 18) caused by a small increase in *f*_*s*_.

Just as the economy of growth is deeply connected to protein allocation, so is growth control. For *A*_*j*_ and *A*_*x*_, this connection is clear from Eqs. 54, 55, respectively. For *A*_*ρ*_, we first note that it relates to optimal marginal values via Eq. 30,

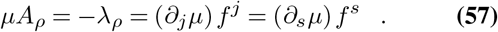

At optimality, the summands on the RHS are zero for the ribosome and for enzymatic reactions (∂_*j*_*μf*_*j*_ = 0 for *j* = r, *e*), and the summation over *j* can thus be restricted to only transporters *s*. Thus, at optimality, the absolute change in optimal growth rate caused by increasing *ρ, μA*_*ρ*_, is equal to the summed marginal effects of transport fluxes on the growth rate, ∂_*s*_*μ*, weighted by the flux fractions *f* ^*s*^ themselves. To see the full connection between *A*_*ρ*_ and protein allocation, we insert Eq. 43 into Eq. 57 to obtain

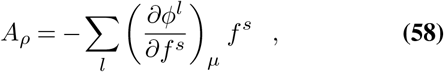

where we sum over *s*. This equation shows that the proportional effect on the optimal growth rate that is exerted by a marginal increase in *ρ, A*_*ρ*_, equals the combined marginal effects of transport fluxes *f* ^*s*^ on proteome allocation fractions, weighted by the transport fluxes themselves.

## Discussion

Modeling frameworks that are essentially linear, such as FBA and RBA, are typically analyzed numerically, as the efficiency of linear programming facilitates fast solutions even for genome-scale models (7, 8, 11). In contrast, the construction and solution of genome-scale non-linear models faces two major obstacles, both intimately linked to the kinetic rate laws. First, experimental estimates for the required kinetic parameters – *k*_cat_ and *K*_m_ values in the simplest case of generalized Michaelis-Menten kinetics – are lacking for most reactions (40). This problem can be alleviated by using parameter estimates from artificial intelligence approaches (41–43). Second, the non-linearity of enzymatic rate laws makes numerical optimizations much more difficult than for linear systems, explaining why existing studies have been limited to models with only a handful of reactions (6, 15–19). Numerical optimization is particularly problematic for models with redundant pathways, where the optimization problem is non-convex (20).

The succinct mathematical formulation for modeling balanced cellular growth developed in this paper helps to address both problems. On the one hand, the reduction of the problem description to a minimal number of independent variables – the flux fractions – reduces the dimensionality of the search space, and can thus accelerate numerical approaches to find optimal states. On the other hand, this formulation allowed us to identify necessary conditions for states of maximal growth rate. These conditions are local for each reaction, i.e., they do not require complete knowledge of the cellular reaction network and its kinetics. In particular, these local balance equations provide a necessary condition for the activity of each individual reaction at optimality: for the corresponding flux to be nonzero, the terms in parentheses in Eqs. 36-38 have to be zero. Our analytical approach thus provides a tool for a deeper understanding of the general principles shaping optimal cellular resource allocation, even when specific optimal states are not explicitly known.

The concise formulation also helped in the interpretation of the optimality conditions from the perspectives of economy and control theory. The marginal change in growth rate induced by each flux change is seen as the flux’s marginal economic value, while the growth adaptation coefficient of each model parameter or external concentration is the change in the optimal growth rate induced by a marginal change in this parameter. The close correspondence between the mathematical expressions obtained in both perspectives helps to clarify the mathematical and conceptual links between these usually separate fields of study, including the extension of previous results of metabolic control analysis (MCA), developed for ad-hoc objectives in static sub-networks, to the holistic problem of cellular growth in GBA models. In MCA, one typically treats enzyme concentrations as control variables and studies how small changes to them affect reactant concentrations and fluxes. Here, all these variables are not only connected, but are uniquely determined by the flux fraction vector **f**. Moreover, the growth rate *μ* itself is explicitly connected to **f** through the growth function (Eq. 16). Through these connections, we can quantify the sensitivity of the cellular growth rate, and hence approximately of organismal fitness, to changes in the control variables ***π***, something not possible in the usual MCA framework (9, 10). The growth adaptation coefficients provide explicit expressions for the effects on growth rate caused by small changes in control variables at optimality. Due to the close relationship between growth rate and fitness, these estimates could be used to interpret and predict evolutionary changes in these variables.

A closely related nonlinear cellular modeling approach accounts for the different amino acid compositions of individual proteins by including “personalized” ribosome reactions for each protein (44–46). In contrast to GBA, this type of model cannot be simplified using flux fractions **f**, as it requires a mathematical formulation that includes explicit variables for metabolite concentrations. Experimental data for *E. coli* (47) indicates that the 20 amino acid content into its total proteome changes very little over 22 highly distinct growth environments (mean coefficient of variation = 2.46%, maximal CV = 7.55%, see Table S1), suggesting that – at least globally – different protein compositions are likely not a major factor driving significant changes in the optimal cellular state. Thus, a unique ribosome reaction with fixed column *M*_r_ is a realistic assumption over all these growth conditions. Further study is necessary to identify whether the different compositions of individual proteins may cause significant changes in their allocation across environments.

All analytical results in this study were derived exclusively from the growth constraints assumed in GBA models: mass conservation in balanced growth, reaction kinetics, cellular density, and non-negative concentrations. For the analysis of optimal growth, we encoded all corresponding information into a single Lagrangian function, parameterized in terms of the constraints. We formulated the problem with the flux fractions **f** as the only free variables, and used KKT conditions to obtain the necessary conditions for optimal growth states. Through these conditions, the marginal protein allocation emerges as the natural underlying currency in the cell economy; this relationship has frequently been asserted (33–35), but is derived here entirely from first principles.

The KKT framework provides a straight-forward way to incorporate new constraints, analogous to how physical theories using the Lagrangian formalism account for additional forces by adding corresponding functions and Lagrange multipliers into the Lagrangian. A re-derivation of the KKT conditions will then result in an extended set of balance equations. Among the potential extra physiological constraints, one might consider also phenomenological constraints such as the recently reported relationship between the cellular surface/volume ratio and the growth rate (48).

One fundamental physiological limitation that could be included in this way but is not considered explicitly here is the diffusion limit of molecules within cellular compartments. This limit links density and kinetic constraints. A higher dry mass density increases the “crowding effect” within cells (25), which entails a lower diffusion rate and by consequence a longer time for reactants to find their catalysts; this effect can be modeled directly by including a corresponding dependence in the Michaelis constants *K*_m_. A study on the crowding effects of all cellular concentrations – including those of small molecules – found that the observed *E. coli* dry mass density is in the range expected if evolution had optimized the cellular density for maximal growth rate (30). In this sense, a fixed density constraint on all molecules, as considered here, may be seen as a simplifying approximation, justified by the observed constancy of cellular buoyant and dry mass densities across different growth conditions (24, 48), with the exception only of large changes in environmental osmolarities (25).

The Lagrangian formalism described here also allows a direct generalization of the theory to other objective functions, i.e., other measures of fitness at balanced growth. This can be done by incorporating a new objective function *F* (**f**) and adding a new constraint for the growth rate via *λ*_*μ*_(*μ* − *μ*_0_), where *μ* is determined by the growth function, *λ*_*μ*_ is the corresponding KKT multiplier, and *μ*_0_ is the constrained growth rate given now as an input.

An important step toward a more general theory of cellular growth would be to extend the GM theory to changing environments, and to derive similar analytical conditions for time-dependent optimal cellular states **f** (*t*). In this situation, fitness is determined by the total growth in a given period of time, so the objective function becomes the integral of the specific growth rate (17, 49), under the same constraints as discussed here. This dynamical extension of the theory might also consider the biologically important scenario of cyclical environments, such as feast-famine cycles of the gut microbiome (50) or day-night cycles of photosynthetic microbes (19).

In sum, the concise mathematical formulation of the growth optimization problem developed here provides a powerful toolbox for the analysis and solution of mechanistic descriptions of optimal cellular physiology and growth. It thereby opens a path toward a fundamental understanding of organizing principles of biological cells. While biological systems will never be fully optimal, optimal states provide an extremely useful null model for the action of natural selection.

## Methods

Let us consider a Lagrangian including inequality constraints on metabolite concentrations,

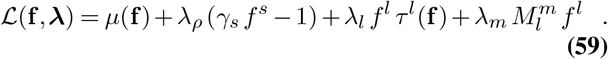

The necessary KKT conditions are the same as in the main text, plus extra conditions for the constraints on metabolite concentrations, including

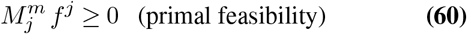

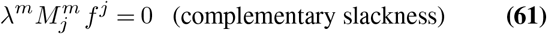

The last equation corresponds to *λ*^*m*^*b*^*m*^ = 0, but since cells in optimal states will only express those proteins that are actually needed to catalyze reactions, we can be even more restrictive and require a stronger version of the last equation:

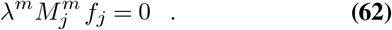

This equation encodes the following: if *λ*^*m*^ ≠ 0, the metabolite *m* is inactive (*c*^*m*^ = 0), and then all of the reactions *j* connected to it (*j* such that 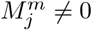) must also be inactive (*f*_*j*_ = 0). We also note that because the turnover times ***τ*** differ from zero, the complementary slackness of reaction *j* (Eq. 27) is equivalent to

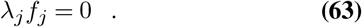

Now we solve the necessary equality conditions, first considering the stationarity

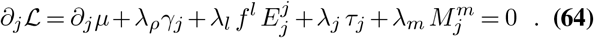

By considering the stronger version of the complementary slackness for reactions (Eq. 63), we cancel the sum on *λ*_*l*_, resulting in

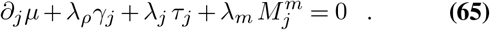

Now we multiply (element-wise) by *f*_*j*_

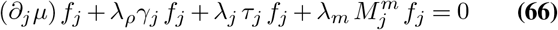

and consider the stronger version of the complementary slackness (Eq. 62) on metabolites, and the complementary slackness on reactions (Eq. 27), resulting in Eq. 29. Summing over all *j* and using the density constraint (Eq. 18) results in Eq. 30. Now the complete expression for *λ*_*ρ*_ in terms of **f** is given by substituting the expression for the growth rate derivatives (Eq. 31),

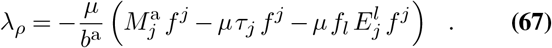

The first term in parentheses equals *b*^a^ according to Eq. 15, and the third equals −*b*^a^ according to Eq. 14, so that both terms cancel each other. This results exactly in Eq. 32.

Finally, combining Eqs. 29, 32, we obtain a general form for the balance equations,

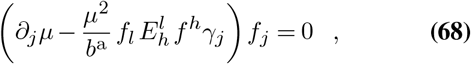

where both indices *l* and *h* are used to indicate summation over all reactions. Using Eq. 31 for the derivative, we have

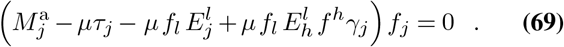

We use Eqs. 9,10,11 to express this in terms of **v, c**,

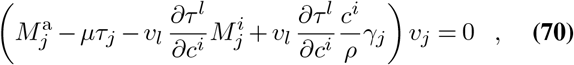

Using *v*_*l*_ = *p*_*l*_*/τ*_*l*_ from Eq. 4, we obtain

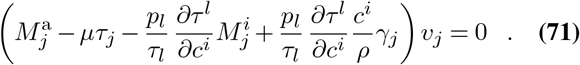

By multiplying out *v*_*j*_, we get Eq. 39.

## ACKNOWLEDGEMENTS

We thank Stefan Müller for discussions about KKT conditions, Alexander Kroll for verifying early calculations, and Xiao-Pan Hu for providing data in Table S1. This work was funded by a Volkswagenstiftung “Life?” grant to MJL, and by the German Research Foundation through grant CRC 1310 to MJL, and, under Germany’s Excellence Strategy, grant EXC 2048/1 (Project ID: 390686111) to OE and MJL.

## Supplementary texts

### Rate laws and kinetic parameters

For simplicity, it may be convenient to assume that each component *τ* ^*j*^ in the vector of turnover times ***τ*** has a general functional form that depends only on a set 𝕂 of kinetic parameters relating reactions *j* with metabolites *m* and external reactants *n*. The simplest general rate law would be the irreversible Michaelis-Menten kinetics, which for a reaction *l* determines

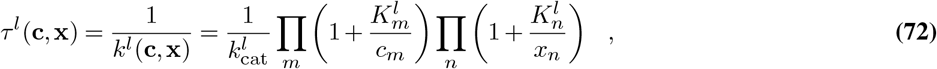

where the kinetic parameters are the corresponding turnover number 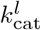 at, and Michaelis constants 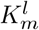 for metabolites *m*, and 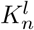 for external reactants *n*. We may consider that all metabolites *m* and external reactants *n* that are not substrates in reaction *l* have corresponding Michaelis constants equal to zero, so the above equation doesn’t depend on their respective concentrations. Note only transport reactions *s* depend on external concentrations **x**, so 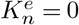 for all *e, n*, and 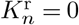 for all *n*.

In that case of all reactions following the irreversible Michaelis-Menten kinetics (72), the (direct) elasticity 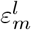 with respect to a metabolite *m* is

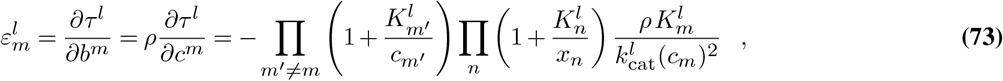

where *m*^*′*^ are metabolites different than *m*. The corresponding indirect elasticity 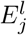 with respect to a reaction *j* is

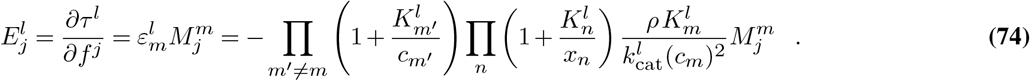

where we note 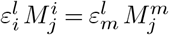 since here 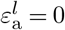 (the Michaelis-Menten kinetics (72) doesn’t depend on the total protein concentration *c*^a^). When *j* = *e* is an enzymatic reaction, the scaling of 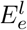 by the respective −*f* ^*e*^ and *τ* ^*l*^ provides the corresponding control coefficient (Eq. 50)

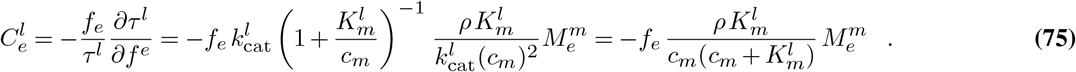

Note that in this case the control coefficients don’t depend on the turnover numbers *k*_cat_, only on Michaelis constants of metabolites *K*_*m*_.

A more realistic example would be some generalized kinetics such as the “convenience kinetics” proposed in Ref. (51), which may also depend on other parameters such as Hill coefficients, inhibitor constants, and stoichiometric coefficients, so these may also be necessary to determine the set of kinetic parameters 𝕂, and by consequence, the model uniquely. With known rate laws, a model is also uniquely determined by the corresponding triple (**M**, 𝕂, *ρ*).

### Mass balance and the stoichiometric matrix S

For a stoichiometric matrix **S** including all reactions and reactants (internal and external ones), and the corresponding vector ***w*** of molecular masses (also known as molecular weights) of reactants, mass conservation within reactions implies

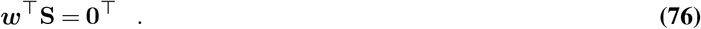

Note that, therefore, the vector of molecular masses must be in the left-null space of **S**.

If we restrict the stoichiometric matrix to internal reactants *i*, then the internal product of ***w***^⊤^ with the columns of this new matrix with entries 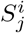 is nonzero only for transporters

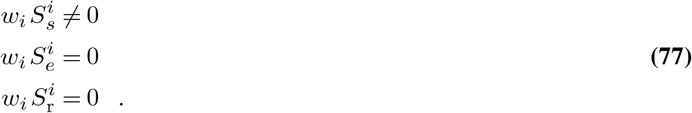

Given that **M** is a mass-scaled version of **S**, these relations are equivalent to Eqs. 2 in the main text; only transporters are capable of increasing or decreasing mass within the model.

We note that our considerations about mass conservation presuppose that all reactants also appear explicitly in the model (i.e. they have a corresponding row in **S**). If some reactants (e.g. water or protons) are omitted from the model for convenience, mass balance is not satisfied. In fact, many models in the literature do ignore some reactants, in particular water; this needs attention in realistic models where water is not only a medium but also a reactant in many biochemical reactions.

### Examples of GBA models and R code for numerical optimization

We present here 4 simple examples of GBA models. We assume for all models a simple irreversible Michaelis-Menten kinetics (72), so in each model turnover times ***τ*** are uniquely defined by a matrix **K** of Michaelis constants and a vector **k**_cat_ of turnover numbers for each reaction. Each row of **K** corresponds to a reactant, and each column to a reaction, matching the order in the matrix **M** (the entries for external reactants are separated by a horizontal line). See the corresponding “.ods” files for a more detailed description of the models, including labels for reactions and reactants and different growth conditions (i.e. external concentrations) for testing the model. Numerical simulations are done by running the R code in the file “GBA.R”, with the variable “modelname” set to the desired model (e.g. modelname ← “A”). Here **K** and *ρ* are in units of *g/L*, and *k*_cat_ in units of *h*^−1^ (resulting from product mass per protein mass per *h*). The code exports the results as a corresponding pdf file with relevant plots for visualization, and a csv file with the numerical values for the optimal cell state at different growth conditions.

Figure (5) presents the schematics and the corresponding parameters (**M, K, k**_cat_, *ρ*) of each model. Metabolites are indicated with circles, and total protein with a rounded square. The numbers labeling reactants and reactions match the corresponding order of rows and columns in **M**, with the last row corresponding to total protein and last column the ribosome reaction by default. All parameters are arbitrary, with the exception of the ribosome reaction where we use 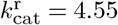 and 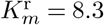 for its main substrate, based on the estimations for *E. coli* in Ref.(4), and *ρ* = 340 based on the measured *E. coli* dry mass density (52).

**Fig. 5.**
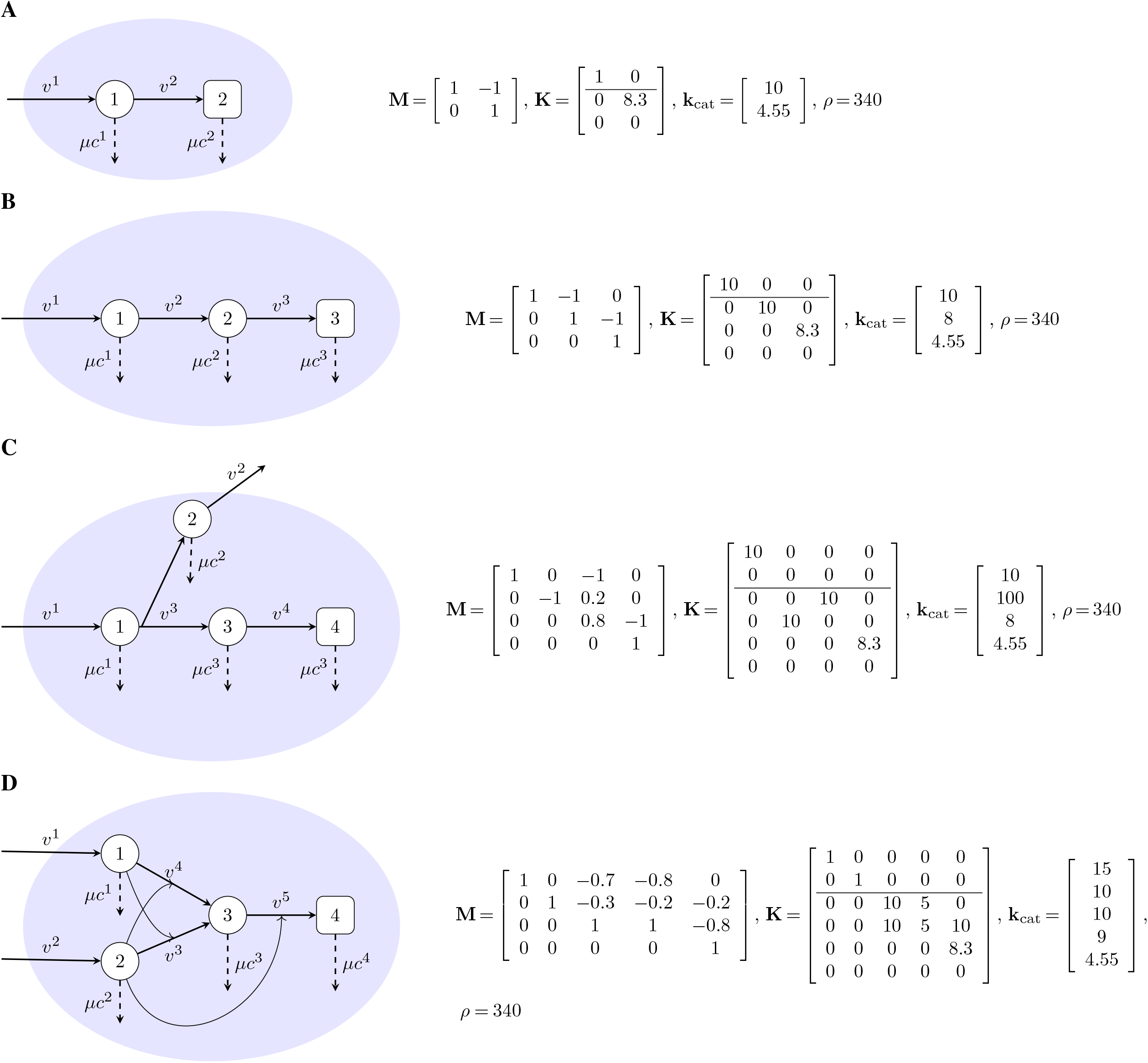
Schematics and parameters defining each model. For more details see the corresponding ods files in Dataset S1.

Models A and B have the simplest model structures (a linear pathway) for two and three reactions, respectively. Model C has a second transport reaction excreting metabolite “2”. Model “D” has two redundant reactions (“3” and “4”), of which only one is active at optimal growth (see optimization results).

### The dependence of *λ*_*ρ*_ on transporters

Equation 29 involves the sums 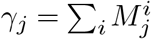 of each column *j* in **M**, which differ from zero only for transporters (Eq. 2). This means Eq. 29 can be separated into distinct equations for *j* = r, *e, s*

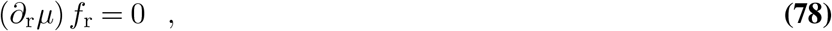

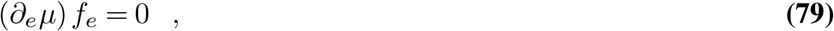

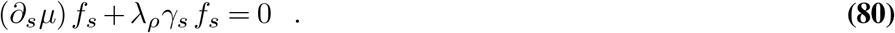

From equations (78,79), we see that, at optimality, the summation in Eq. 30 can be simplified to a summation over *s* only

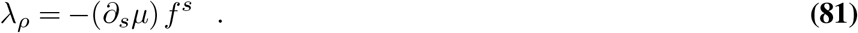

Substituting now the partial derivative given by Eq. 31, we obtain

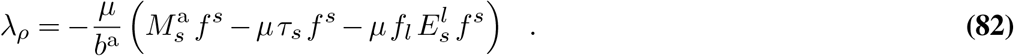

The first summand in the parenthesis equals to zero, since transporters do not produce protein 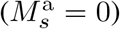, and the second summand can be expressed in terms of protein fractions *ϕ*^*s*^ = *p*^*s*^*/c*^a^ = *μτ* ^*s*^ *f* ^*s*^*/b*^a^, resulting in

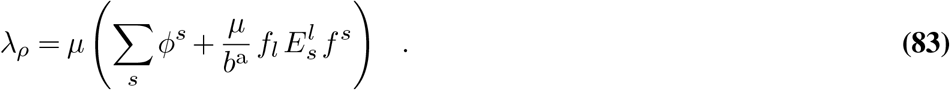

### Optimal enzyme concentrations and control coefficients

Equation 50 shares a striking analogy with an optimality condition for metabolic systems, established in (36): in an optimal metabolic state, maximizing a pathway flux at a limited total enzyme amount, the enzyme levels must be proportional to the flux control coefficients. This previous result reflects the assumption that the cell trades a cost (the sum of enzyme levels) against a benefit (the pathway flux), and that in an optimal state, the marginal cost and benefit, for any small change of enzyme levels, must be the same. Equation 50, for optimal growth states, has a similar interpretation, but without an explicit benefit function for fluxes. Here, instead, if an enzyme level in reaction A has an indirect effect on a flux in reaction B and makes reaction B proceed more efficiently, then catalyzing enzyme for reaction B can be saved (and resources be redistributed to increase growth). Hence, we now have a trade-off between a cost (of investing enzyme in A) and an “opportunity benefit” (enzyme saved in reaction B). In the equation, an enzyme of interest (in reaction A) is described by *p*_*e*_, its effect on all fluxes in the network (reactions B) is described by the control coefficients 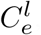, and the catalysts of these reactions are represented by *p*_*l*_. By summing all marginal “opportunity benefits” 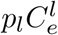, we obtain the equivalent to the marginal flux benefit in (36).

From the similar Eq. 44 in the main text, we can infer that at optimality, no two active enzymes can realistically catalyze the exact same reaction (i.e., have identical columns in **M**). If this were the case, then both turnover times *τ*_*e*_ would have to be exactly the same, since the marginal opportunity would be identical for both enzymes (see Eq. 24). This condition is highly unlikely to hold in realistic models, since real enzymes will always have different physical properties and therefore different kinetics. Thus, if several isoenzymes catalyze a particular reaction in a model, only one of these reactions will be active at optimality. The previous argument can be generalized to any two linear combinations of reactions in the model, and as a consequence, the submatrix resulting from restricting **M** to active reactions must have full column rank at optimal growth (see Refs. (4, 21, 22)). Note that in reality enzymes with very similar marginal values may still be expressed together due to the little selection pressure towards one of them. See model “D” on SI text “Examples of GBA models and R code for numerical optimization” for an example of a model with redundant reactions.

### Amino acid composition of the *E. coli* proteome

Table S1 presents the amino acid frequency in the *E. coli* proteome, calculated from protein sequence (retrieved from genome annotation of *E. coli* NC_000913.3 from RefSeq (53)) and weighted by protein abundance measured in Schmidt et al. (47) under various growth conditions.

**Table S1.**
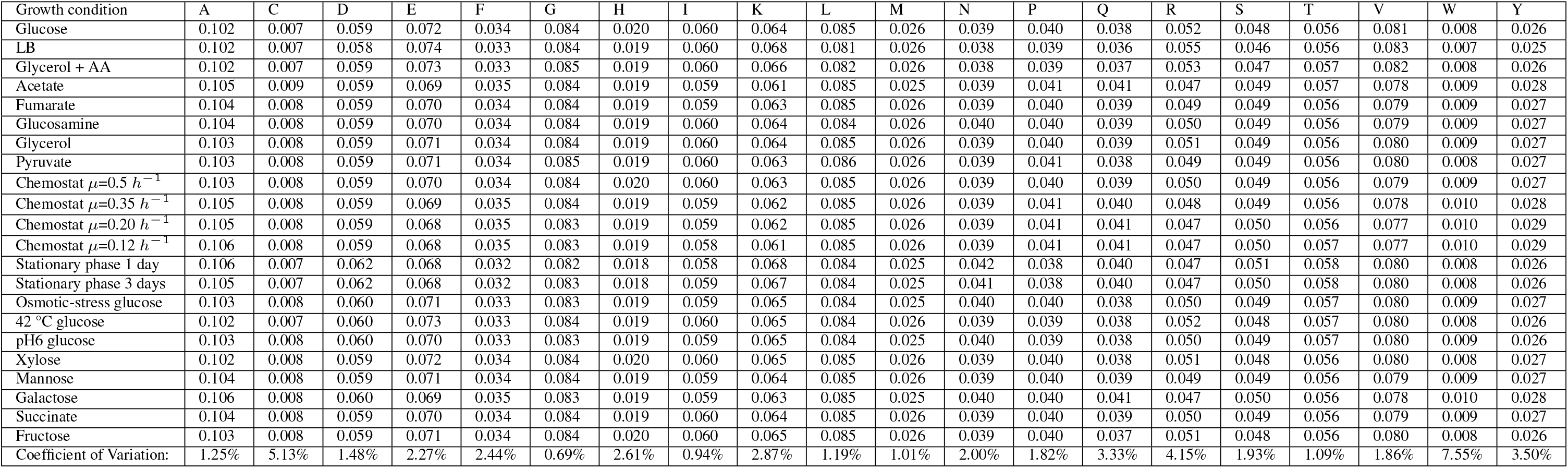
Amino acid frequency in the *E. coli* proteome at various growth conditions.

### SI_files.zip

Zip file containing files for numerical optimization, as described in the SI text “Examples of GBA models and R code for numerical optimization”.

