## Supplementary material for "Growth Mechanics: General principles of optimal cellular resource allocation in balanced growth": SI files: Exportcsv.R

```
# exporting file with the optimal states ci, tau, mu, v, p 

opt_state <- matrix(rep(0,(nx+2+ni+nj+nj+nj)*n_conditions),nrow = n_conditions)
for (cond in 1:n_conditions) {
  
  rho <- rho_cond[cond]
  
  x  <- x_cond[,cond]
  
  f <- f_opt[cond,]
  
  opt_state[cond,] <- c(conv[cond],mu(f),x,ci(f),tau(ci(f)),v(f),p(f))
}

colnames(opt_state) <- c("convergence","mu",reactant,paste("tau",reaction),
                         paste("v",reaction),paste("p",reaction))

# export results
write.csv(opt_state, file = paste("Model ",modelname,
                            ", mean time (",mean_time,"s) results.csv",sep=""))
```
