## Supplementary material for "Growth Mechanics: General principles of optimal cellular resource allocation in balanced growth": SI files: f0.R

```
# Initial value subproblem: linear optimization to find maximal ribosome flux
# fraction f^r, with a minimal production of each metabolite. The constraints 
# are mass conservation (M*f = b) and surface flux balance (sM*f = 1)

# minimal biomass fractions b

min_b <- 1/ni/10

# defining parameters

objective.fn <- c(rep(0,nj-1),1)
const.mat <- rbind(M[-ni,],sM)
const.dir <- c(rep(">=",ni-1),"=")
const.rhs <- c(rep(min_b,ni-1), 1)

# solving model

lp.solution <- lp("max", objective.fn, const.mat, const.dir, const.rhs)

# solution to the linear optimization

f0 <- lp.solution$solution

# stops if linear optimization didn't find solution

if (sum(f0) == 0)  stop("no feasible solution found for initial f0")

# stops if solution is not a valid state for the nonlinear problem due to
# negative growth rate or concentrations

rho <- rho_cond[1]

x  <- x_cond[,1]

if (TRUE %in% c(mu(f0) < 0, p(f0) < 0, ci(f0) < 0)) stop("initial f0 is not a 
                                    valid state for the nonlinear optimization")
```
