## Supplementary material for "Growth Mechanics: General principles of optimal cellular resource allocation in balanced growth": SI files: GBA.R

```
# GBA model solver

# Clear variables
rm(list=ls(all=TRUE))

require('rstudioapi') 
require('readODS')
require('nloptr')
require('Matrix')
require('lpSolve')

# Sets working directory to source file location ###############################

directory <- dirname(getActiveDocumentContext()$path)

setwd(directory) 

# Model name here ##############################################################

modelname <- "D"

# Reads model saved as .ods file ###############################################

source("Readmodelods.R")

# Definitions ##################################################################

# index for external reactants
n <- 1: dim(x_cond)[1]

# number of external reactants
nx <- dim(x_cond)[1]

# number of growth conditions
n_conditions <- dim(x_cond)[2]

# names of internal reactants
i_reactant <- reactant[-n]

# number of internal reactants
ni <- dim(M)[1]

# number of reactions
nj <- dim(M)[2]

# the sum of each M column 
sM <- colSums(M)

# delete numerical artifacts
sM[abs(sM) < 1e-10] <- 0

# indexes for reactions: s (transport), e (enzymatic), and ribosome r 

e <- c(1:(nj-1))[sM[1:(nj-1)] == 0]  

s <- c(1:(nj-1))[-e]

r <- nj   # (the ribosome is by default the last reaction)

# indexes: m (metabolite), a (all proteins)

m <- 1:(ni-1)

a <- ni   # (the last row by default corresponds to all proteins)

# number of transport reactions
ns <- length(s)

# tau (irreversible Michaelis-Menten kinetics) #################################
tau <- function(c){
  
  # all concentrations x,c
  xc <- c(x, c) 
  
  tauj <- rep(0,nj)
  
  for (j in 1:nj) {
    
    tauj[j] <- as.numeric(prod(1 + K[,j]/xc)/kcat[j]) 
    
  }
  
  return(tauj)
}

# derivative with respect to c
dtau <- function(c){
  
  # all concentrations x,c
  xc <- c(x, c) 
  
  ditauj <- matrix(rep(0,nj*ni),ncol=ni)
  
  for (j in 1:nj) {
    
    for (i2 in 1:ni) {
      
      y <- i2 + nx # position of i in terms of all reactants xc
      
      ditauj[j,i2] <- -as.numeric((K[y,j]/(c[i2]^2))*prod(1 + 
                                                            K[-y,j]/xc[-y])/kcat[j]) 
      
    }
    
  }
  
  return(ditauj)
  
}

# Optimization #################################################################

source("Optimization.R") 

# Showing results ##############################################################

# Exporting csv file with results #######

source("Exportcsv.R")

# Plots #################################

source("Plots.R")
```
