## Supplementary material for "Growth Mechanics: General principles of optimal cellular resource allocation in balanced growth": SI files: Optimization.R

```
# Growth optimization

# Functions ####################################################################

# mu ############## (growth rate)
mu <- function(f) as.numeric(M[a,r]*f[r]/(tau(ci(f))%*%f)) 

# negative mu (for minimization)
negative_mu <- function(f) -as.numeric(M[a,r]*f[r]/(tau(ci(f))%*%f)) 

# fluxes
v <- function(f) as.vector(mu(f))*rho*f

# protein concentrations
p <- function(f) tau(ci(f))*v(f)

# internal concentrations "i" of metabolites "m" and total protein "a"
ci <- function(f) rho*M%*%f

# biomass fractions
b <- function(f) M%*%f

# Optimization #################################################################

# equality constraints (density constraint)

g <- function(f) sM%*%f - 1

# inequality constraints (non negativity of concentrations). Here we assume ####
# mu ~ 1 for a faster calculation of p, instead of using the function p(f) #####

h <- function(f) c(ci(f), rho*tau(ci(f))*f)

# Gradient of negative_mu ######################################################

negative_dmu <- function(f) -((mu(f)^2)/b(f)[a])*(M[a,]/mu(f) - 
                                      t(f)%*%(rho*dtau(ci(f))%*%M) - tau(ci(f)))

# Jacobian of equality constraint ##############################################

dg	<- function(f) sM

# Jacobian of inequality constraints ###########################################

# rows = c1, c2,...,p1,p2... , columns = d/df1, d/df2, ...
dh	<- function(f) rho*rbind(M,rho*(dtau(ci(f))%*%M*matrix(rep(f,nj),nrow=nj)) + 
                              diag(nj)*tau(ci(f)) )

# Finds initial f0 #############################################################

source('f0.R')

# starts loop for the optimization at each environmental condition #############
f_opt  <- matrix(rep(0,nj*n_conditions),ncol=nj)
mu_opt <- rep(0,n_conditions)
otime  <- rep(0,n_conditions)
conv   <- rep(0,n_conditions)
iter   <- rep(0,n_conditions)
for (cond in 1:n_conditions) {
  
  rho <- rho_cond[cond]
  
  x  <- x_cond[,cond]
  
  # Upper bounds 
  upper_f <- rep(2,nj)
  
  # lower bounds 
  lower_f <- rep(-2,nj)
  
  # Optimization
  solver      <- "LBFGS" # Local solvers: "COBYLA", "LBFGS", "MMA", or "SLSQP"
  
  # measuring the total optimization time
  st <- system.time({
    
    # optimization (by default the function auglag minimizes, so we use 
    # negative_mu instead of mu as the objective function)
    
    res <- auglag(f0, fn = negative_mu, gr = negative_dmu, lower = lower_f, 
                  upper = upper_f, heq = g, heqjac = dg, hin = h,hinjac = dh, 
                  localsolver = c(solver), localtol = 1e-5,
                  control = list(maxeval = 1000))
    
  })
  
  # solution
  f_opt[cond,] <- res$par
  
  # optimal mu
  mu_opt[cond] <- mu(f_opt[cond,])
  
  # convergence (codes: "-1" means optimization problem, "5" means optimization 
  # stop because maxeval, "4" means optimization stopped because xtol_rel or 
  # xtol_abs (above) was reached)
    conv[cond] <- res$convergence
  
  # optimization time
  otime[cond] <- signif(st[[1]], digits= 4)
  
  # number of iterations
  iter[cond] <- res$iter  
  
  print(paste("optimization: ",cond, "/", n_conditions,", optimization time: ",
              otime[cond]," s, convergence: ", conv[cond],", growth rate: ",
              signif(mu_opt[cond], digits= 3), sep=""))
  
  # next initial value
  f0 <- res$par
  
}

# produces c_opt
c_opt <- matrix(rep(0,ni*n_conditions),ncol=ni)
for (cond in 1:n_conditions) c_opt[cond,] <- ci(f_opt[cond,])

# mean optimization time
mean_time <- signif(mean(otime), digits= 3)
```
