## Supplementary material for "Growth Mechanics: General principles of optimal cellular resource allocation in balanced growth": SI files: Plots.R

```
# plot results #######################################################################################################

pdf(paste("Model ",modelname,", mean time (",mean_time,"s) results.pdf",sep=""),title="Optimization results",
    width=10,height=5)
par(mfrow=c(1,2))

# graphic
coloraxis <- "darkgrey"
colorlab <- "black"

# Growth rate vs. log10(first external concentration) ################################################################

mu_opt <- opt_state[,"mu"]
cx1    <- log10(opt_state[,reactant[1]])

minx1 <- floor(min(cx1))
maxx1 <- ceiling(max(cx1)*1.1)
maxmu <- max(mu_opt)*1.1

plot(cx1,mu_opt,pch=16, mgp=c(2, 0.5, 0), col.lab = colorlab, frame.plot=F, col= 1,xlim=c(minx1,maxx1),ylim=c(0,maxmu), xaxt = "n",
     yaxt = "n", ylab=bquote("Growth rate"~ mu ~ (h^-1)), xlab=paste("Concentration",reactant[1],"log10(g/L)"), yaxs="i", xaxs="i")
axis(1, at=seq(minx1,maxx1,1), mgp=c(3, 0.4, 0), col=coloraxis, col.ticks=coloraxis, col.axis=coloraxis, tck=0.02,las=1)
axis(1, at=seq(minx1,maxx1,1), mgp=c(3, 0.4, 0), col=coloraxis, col.ticks=coloraxis, col.axis=coloraxis, tck=-0.02,las=1)
axis(2, mgp=c(3, 0.4, 0), col=coloraxis, col.ticks=coloraxis, col.axis=coloraxis, tck=0.02,las=1)
axis(2, mgp=c(3, 0.4, 0), col=coloraxis, col.ticks=coloraxis, col.axis=coloraxis, tck=-0.02,las=1)

if (nx > 1) {
  # Growth rate vs. second external concentration ####################################################################
  
  cx2    <- log10(opt_state[,reactant[2]])
  
  minx2 <- floor(min(cx2))
  maxx2 <- ceiling(max(cx2)*1.1)
  
  plot(cx2,mu_opt,pch=16, mgp=c(2, 0.5, 0), col.lab = colorlab, frame.plot=F, col= 1,ylim=c(0,maxmu), xaxt = "n",
       yaxt = "n", ylab=bquote("Growth rate"~ mu ~ (h^-1)), xlab=paste("Concentration",reactant[2],"log10(g/L)"), yaxs="i", xaxs="i")
  axis(1, mgp=c(3, 0.4, 0), col=coloraxis, col.ticks=coloraxis, col.axis=coloraxis, tck=0.02,las=1)
  axis(1, mgp=c(3, 0.4, 0), col=coloraxis, col.ticks=coloraxis, col.axis=coloraxis, tck=-0.02,las=1)
  axis(2, mgp=c(3, 0.4, 0), col=coloraxis, col.ticks=coloraxis, col.axis=coloraxis, tck=0.02,las=1)
  axis(2, mgp=c(3, 0.4, 0), col=coloraxis, col.ticks=coloraxis, col.axis=coloraxis, tck=-0.02,las=1)
  
} else { 
  
  # Growth rate vs. log10(first external concentration) ##############################################################
  
  cx1    <- opt_state[,reactant[1]]
  
  minx1 <- 0
  maxx1 <- ceiling(max(cx1)*1.1)
  
  plot(cx1,mu_opt,pch=16, mgp=c(2, 0.5, 0), col.lab = colorlab, frame.plot=F, col= 1,xlim=c(minx1,maxx1),ylim=c(0,maxmu), xaxt = "n",
       yaxt = "n", ylab=bquote("Growth rate"~ mu ~ (h^-1)), xlab=paste("Concentration",reactant[1],"(g/L)"), yaxs="i", xaxs="i")
  axis(1, mgp=c(3, 0.4, 0), col=coloraxis, col.ticks=coloraxis, col.axis=coloraxis, tck=0.02,las=1)
  axis(1, mgp=c(3, 0.4, 0), col=coloraxis, col.ticks=coloraxis, col.axis=coloraxis, tck=-0.02,las=1)
  axis(2, mgp=c(3, 0.4, 0), col=coloraxis, col.ticks=coloraxis, col.axis=coloraxis, tck=0.02,las=1)
  axis(2, mgp=c(3, 0.4, 0), col=coloraxis, col.ticks=coloraxis, col.axis=coloraxis, tck=-0.02,las=1)
  
}

# Internal reactant concentrations vs.mu #############################################################################

colori <- rainbow(ni)

maxci <- rho 

matplot(mu_opt,c_opt,pch=16, mgp=c(2, 0.5, 0), col.lab = colorlab, frame.plot=F, col= colori,xlim=c(0,maxmu),ylim=c(0,maxci), xaxt = "n",
        yaxt = "n", xlab=bquote("Growth rate"~ mu ~ (h^-1)), ylab=bquote("Internal concentrations " ~ c^i ~ " (g/L)"), yaxs="i", xaxs="i")
axis(1, mgp=c(3, 0.4, 0), col=coloraxis, col.ticks=coloraxis, col.axis=coloraxis, tck=0.02,las=1)
axis(1, mgp=c(3, 0.4, 0), col=coloraxis, col.ticks=coloraxis, col.axis=coloraxis, tck=-0.02,las=1)
axis(2, mgp=c(3, 0.4, 0), col=coloraxis, col.ticks=coloraxis, col.axis=coloraxis, tck=0.02,las=1)
axis(2, mgp=c(3, 0.4, 0), col=coloraxis, col.ticks=coloraxis, col.axis=coloraxis, tck=-0.02,las=1)
legend("topright",i_reactant,col=colori,pch=16,cex=0.7)

# Metabolite concentrations vs. mu ###################################################################################

cm_opt <- c_opt[,-a]

maxcm <- ceiling(max(cm_opt))

matplot(mu_opt,cm_opt,pch=16, mgp=c(2, 0.5, 0), col.lab = colorlab, frame.plot=F, col= colori,xlim=c(0,maxmu),ylim=c(0,maxcm), xaxt = "n",
        yaxt = "n", xlab=bquote("Growth rate"~ mu ~ (h^-1)), ylab=bquote("Metabolite concentrations " ~ c^m ~ " (g/L)"), yaxs="i", xaxs="i")
axis(1, mgp=c(3, 0.4, 0), col=coloraxis, col.ticks=coloraxis, col.axis=coloraxis, tck=0.02,las=1)
axis(1, mgp=c(3, 0.4, 0), col=coloraxis, col.ticks=coloraxis, col.axis=coloraxis, tck=-0.02,las=1)
axis(2, mgp=c(3, 0.4, 0), col=coloraxis, col.ticks=coloraxis, col.axis=coloraxis, tck=0.02,las=1)
axis(2, mgp=c(3, 0.4, 0), col=coloraxis, col.ticks=coloraxis, col.axis=coloraxis, tck=-0.02,las=1)
legend("topleft",i_reactant[-a],col=colori,pch=16,cex=0.7)

# Protein concentrations #############################################################################################

colorj <- 1:nj

if (nj > 8) colorj <- rainbow(nj)

p_opt <- opt_state[,paste("p",reaction)]

maxp <- ceiling(max(p_opt)*1.1)

matplot(mu_opt,p_opt,pch=16, mgp=c(2, 0.5, 0), col.lab = colorlab, frame.plot=F, col= colorj,xlim=c(0,maxmu),ylim=c(0,maxp), xaxt = "n",
        yaxt = "n", xlab=bquote("Growth rate"~ mu ~ (h^-1)), ylab=bquote("Protein concentrations " ~  p ~ " (g/L)"), yaxs="i", xaxs="i")
axis(1, mgp=c(3, 0.4, 0), col=coloraxis, col.ticks=coloraxis, col.axis=coloraxis, tck=0.02,las=1)
axis(1, mgp=c(3, 0.4, 0), col=coloraxis, col.ticks=coloraxis, col.axis=coloraxis, tck=-0.02,las=1)
axis(2, mgp=c(3, 0.4, 0), col=coloraxis, col.ticks=coloraxis, col.axis=coloraxis, tck=0.02,las=1)
axis(2, mgp=c(3, 0.4, 0), col=coloraxis, col.ticks=coloraxis, col.axis=coloraxis, tck=-0.02,las=1)
legend("topright",reaction,col=colorj,pch=16,cex=0.7)

# Protein fractions ##################################################################################################

phi_opt <- p_opt/c_opt[,a]

maxp <- ceiling(max(p_opt)*1.1)

matplot(mu_opt,phi_opt,pch=16, mgp=c(2, 0.5, 0), col.lab = colorlab, frame.plot=F, col= colorj,xlim=c(0,maxmu),ylim=c(0,1), xaxt = "n",
        yaxt = "n", xlab=bquote("Growth rate"~ mu ~ (h^-1)), ylab=bquote("Proteome fractions " ~  phi ), yaxs="i", xaxs="i")
axis(1, mgp=c(3, 0.4, 0), col=coloraxis, col.ticks=coloraxis, col.axis=coloraxis, tck=0.02,las=1)
axis(1, mgp=c(3, 0.4, 0), col=coloraxis, col.ticks=coloraxis, col.axis=coloraxis, tck=-0.02,las=1)
axis(2, mgp=c(3, 0.4, 0), col=coloraxis, col.ticks=coloraxis, col.axis=coloraxis, tck=0.02,las=1)
axis(2, mgp=c(3, 0.4, 0), col=coloraxis, col.ticks=coloraxis, col.axis=coloraxis, tck=-0.02,las=1)
legend("top",reaction,col=colorj,pch=16,cex=0.7)

# Fluxes vs. mu ######################################################################################################

v_opt <- opt_state[,paste("v",reaction)]

maxv <- ceiling(max(v_opt)*1.1)

matplot(mu_opt,v_opt,pch=16, mgp=c(2, 0.5, 0), col.lab = colorlab, frame.plot=F, col= colorj,xlim=c(0,maxmu),ylim=c(0,maxv), xaxt = "n",
        yaxt = "n", xlab=bquote("Growth rate"~ mu ~ (h^-1)), ylab=bquote("Fluxes " ~  v ~ " (g/L/h)"), yaxs="i", xaxs="i")
axis(1, mgp=c(3, 0.4, 0), col=coloraxis, col.ticks=coloraxis, col.axis=coloraxis, tck=0.02,las=1)
axis(1, mgp=c(3, 0.4, 0), col=coloraxis, col.ticks=coloraxis, col.axis=coloraxis, tck=-0.02,las=1)
axis(2, mgp=c(3, 0.4, 0), col=coloraxis, col.ticks=coloraxis, col.axis=coloraxis, tck=0.02,las=1)
axis(2, mgp=c(3, 0.4, 0), col=coloraxis, col.ticks=coloraxis, col.axis=coloraxis, tck=-0.02,las=1)
legend("topleft",reaction,col=colorj,pch=16,cex=0.7)

# flux fractions vs. mu ##############################################################################################

w_opt <- v_opt/(mu_opt*rho_cond)

maxw <- ceiling(max(w_opt))

matplot(mu_opt,w_opt,pch=16, mgp=c(2, 0.5, 0), col.lab = colorlab, frame.plot=F, col= colorj,xlim=c(0,maxmu),ylim=c(0,maxw), xaxt = "n",
        yaxt = "n", xlab=bquote("Growth rate"~ mu ~ (h^-1)), ylab=bquote("Flux fractions " ~  f), yaxs="i", xaxs="i")
axis(1, mgp=c(3, 0.4, 0), col=coloraxis, col.ticks=coloraxis, col.axis=coloraxis, tck=0.02,las=1)
axis(1, mgp=c(3, 0.4, 0), col=coloraxis, col.ticks=coloraxis, col.axis=coloraxis, tck=-0.02,las=1)
axis(2, mgp=c(3, 0.4, 0), col=coloraxis, col.ticks=coloraxis, col.axis=coloraxis, tck=0.02,las=1)
axis(2, mgp=c(3, 0.4, 0), col=coloraxis, col.ticks=coloraxis, col.axis=coloraxis, tck=-0.02,las=1)
legend("topright",reaction,col=colorj,pch=16,cex=0.7)

# Turnover times ####################################################################################################

tau_opt <- opt_state[,paste("tau",reaction)]

maxtau <- ceiling(max(tau_opt))

matplot(mu_opt,tau_opt,pch=16, mgp=c(2, 0.5, 0), col.lab = colorlab, frame.plot=F, col= colorj,xlim=c(0,maxmu),ylim=c(0,maxtau), xaxt = "n",
        yaxt = "n", xlab=bquote("Growth rate"~ mu ~ (h^-1)), ylab=bquote("Turnover times " ~  tau ~ (h)), yaxs="i", xaxs="i")
axis(1, mgp=c(3, 0.4, 0), col=coloraxis, col.ticks=coloraxis, col.axis=coloraxis, tck=0.02,las=1)
axis(1, mgp=c(3, 0.4, 0), col=coloraxis, col.ticks=coloraxis, col.axis=coloraxis, tck=-0.02,las=1)
axis(2, mgp=c(3, 0.4, 0), col=coloraxis, col.ticks=coloraxis, col.axis=coloraxis, tck=0.02,las=1)
axis(2, mgp=c(3, 0.4, 0), col=coloraxis, col.ticks=coloraxis, col.axis=coloraxis, tck=-0.02,las=1)
legend("topright",reaction,col=colorj,pch=16,cex=0.7)

# Turnover frequencies ##############################################################################################

f_opt <- 1/tau_opt

maxf <- ceiling(max(f_opt))

matplot(mu_opt,f_opt,pch=16, mgp=c(2, 0.5, 0), col.lab = colorlab, frame.plot=F, col= colorj,xlim=c(0,maxmu),ylim=c(0,maxf), xaxt = "n",
        yaxt = "n", xlab=bquote("Growth rate"~ mu ~ (h^-1)), ylab=bquote("Apparent turnover numbers " ~  k[app] ~ (h^-1)), yaxs="i", xaxs="i")
axis(1, mgp=c(3, 0.4, 0), col=coloraxis, col.ticks=coloraxis, col.axis=coloraxis, tck=0.02,las=1)
axis(1, mgp=c(3, 0.4, 0), col=coloraxis, col.ticks=coloraxis, col.axis=coloraxis, tck=-0.02,las=1)
axis(2, mgp=c(3, 0.4, 0), col=coloraxis, col.ticks=coloraxis, col.axis=coloraxis, tck=0.02,las=1)
axis(2, mgp=c(3, 0.4, 0), col=coloraxis, col.ticks=coloraxis, col.axis=coloraxis, tck=-0.02,las=1)
legend("topleft",reaction,col=colorj,pch=16,cex=0.7)

dev.off()
```
