## Supplementary material for "Growth Mechanics: General principles of optimal cellular resource allocation in balanced growth": SI files: Readmodelods.R

```
# Loads model saved as .ods file ###############################################

odsfile <- paste(modelname,".ods", sep = "")

nsheets    <- get_num_sheets_in_ods(odsfile)
sheets <- list_ods_sheets(odsfile)

# Position of parameters
posM          <- (1:nsheets)[sheets == "M"]
posK          <- (1:nsheets)[sheets == "K"]
poskcat       <- (1:nsheets)[sheets == "kcat"]
posconditions <- (1:nsheets)[sheets == "conditions"]

# Getting data from sheets

# Mass fraction matrix Mni including external reactants
M <- as.matrix(read_ods(odsfile, sheet= posM)[,-1])

# reaction and reactant names
reaction <- colnames(M)
rownames(M) <- read_ods(odsfile, sheet= posM)[,1]

# Michaelis constant matrix K 
K <- as.matrix(read_ods(odsfile, sheet= posK)[,-1])

# kcat
kcat <- as.numeric(read_ods(odsfile, sheet= poskcat)[,-1])

# Growth condition names
condition <- colnames(read_ods(odsfile, sheet= posconditions))[-1]

# Mass density rho at each condition
rho_cond <- as.numeric(read_ods(odsfile, sheet= posconditions)[1,-1])

# external concentrations at each condition
x_cond  <- as.matrix(read_ods(odsfile, sheet= posconditions)[-1,-1])

reactant <- c(read_ods(odsfile, sheet= posconditions)[-1,1],rownames(M))
```
