## Supplementary figures and images for "Growth Mechanics: General principles of optimal cellular resource allocation in balanced growth"

### Model A, mean time (0.0216s) results.pdf

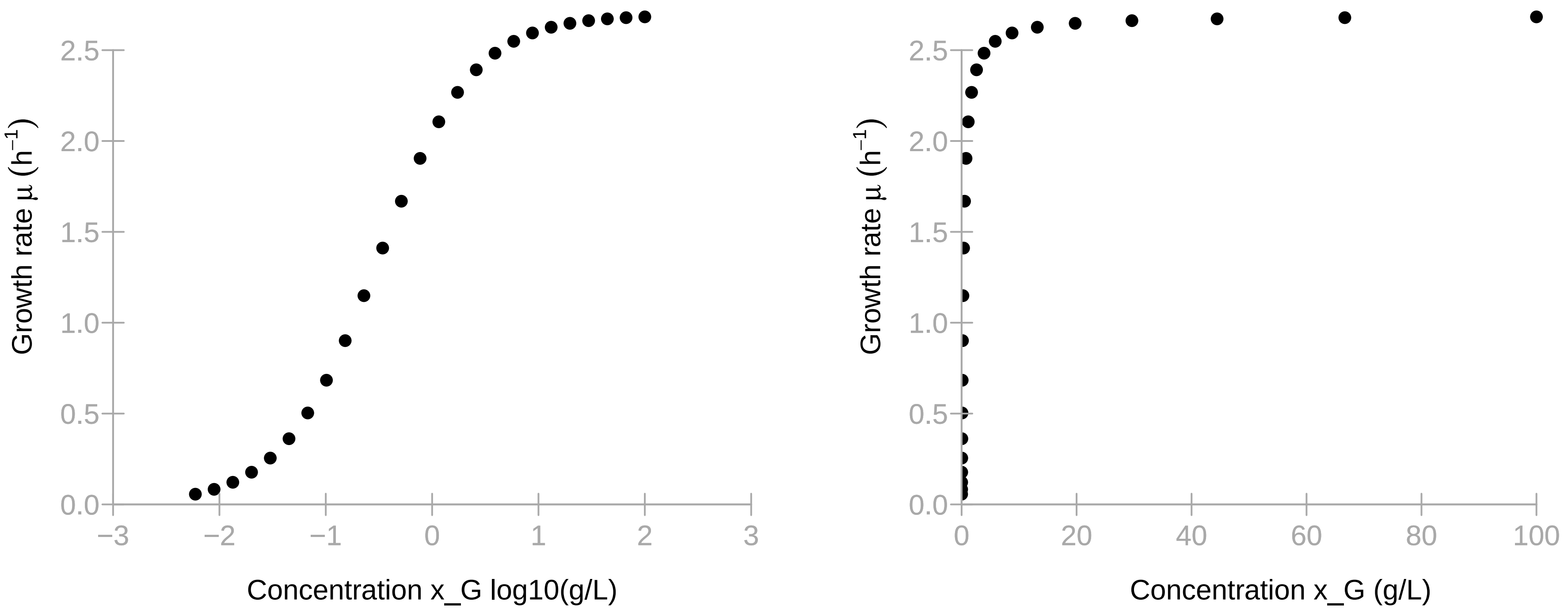

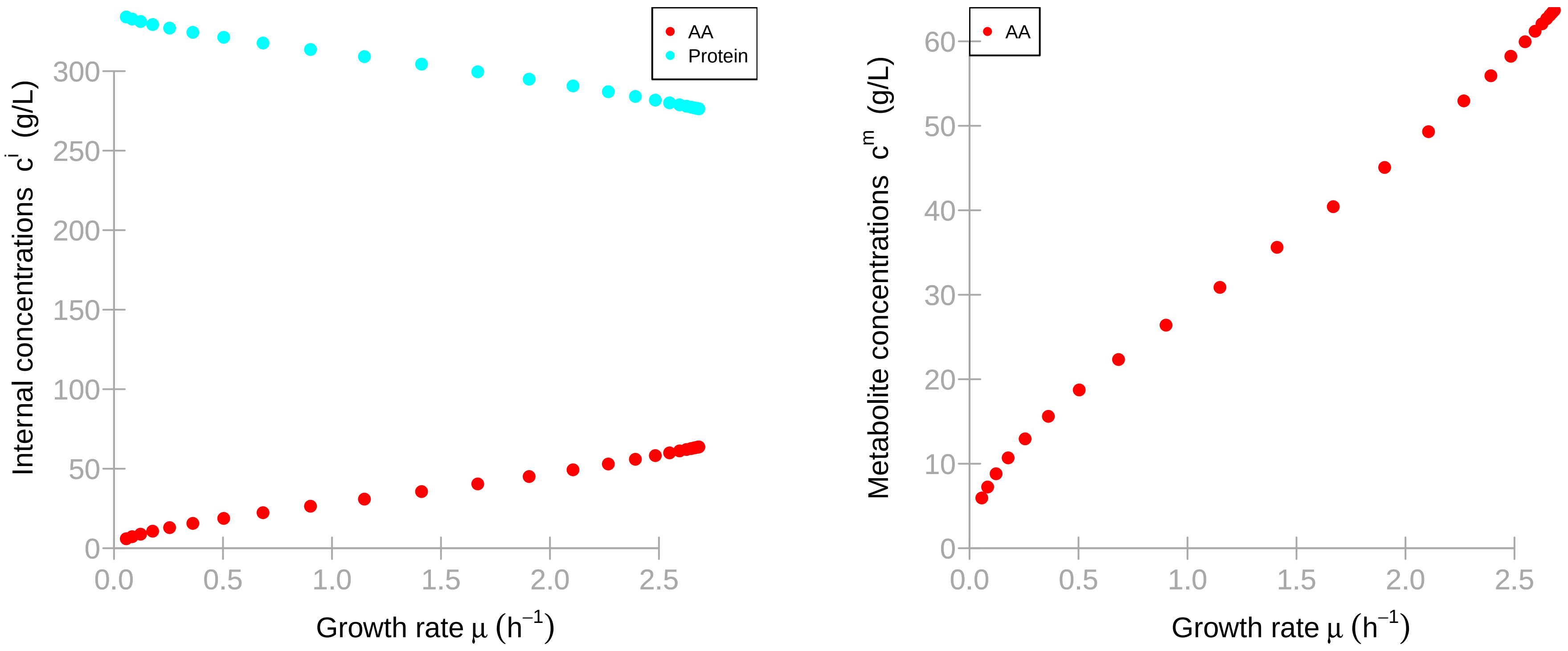

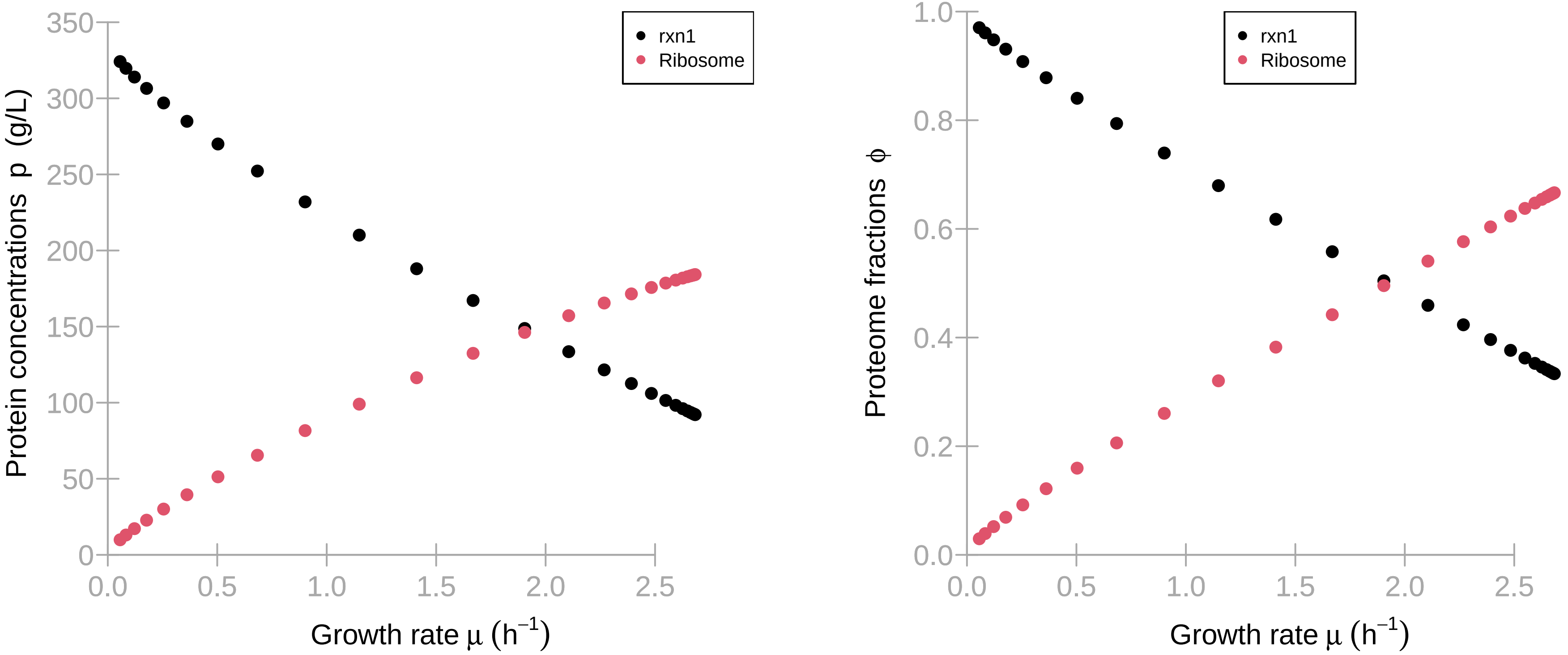

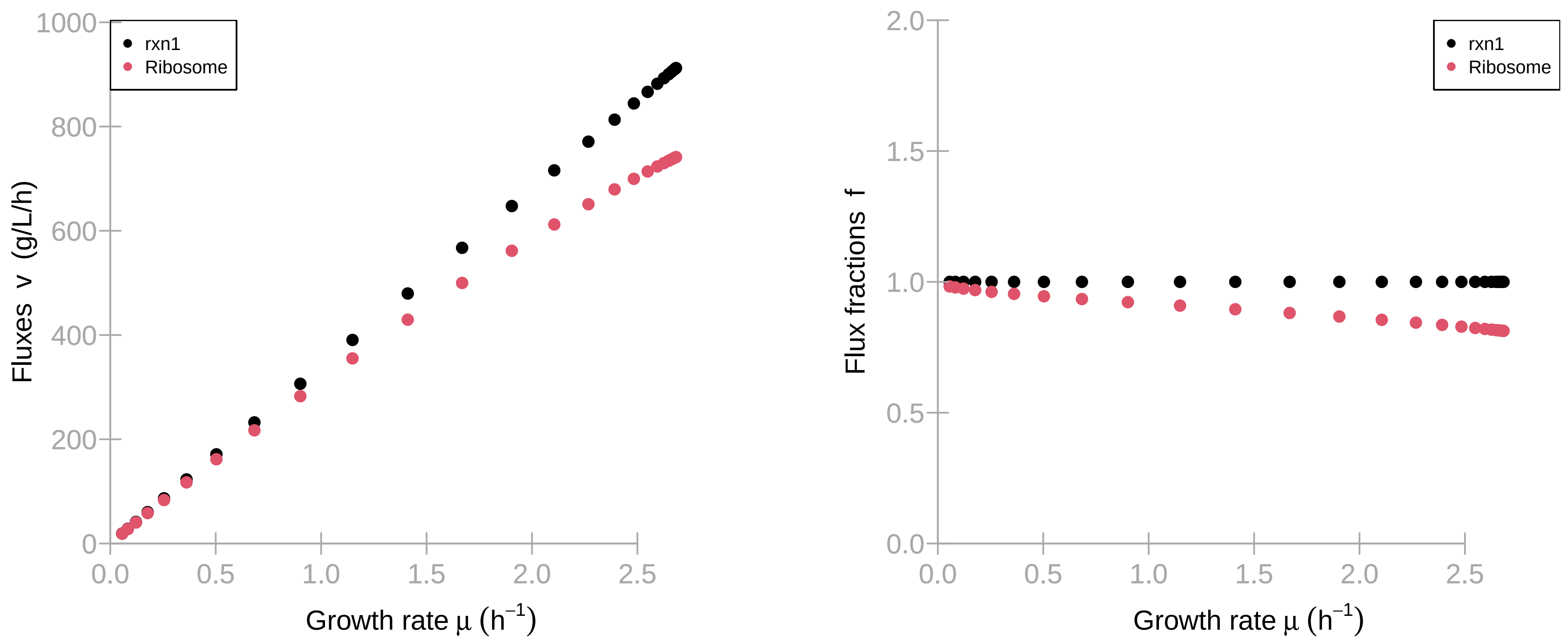

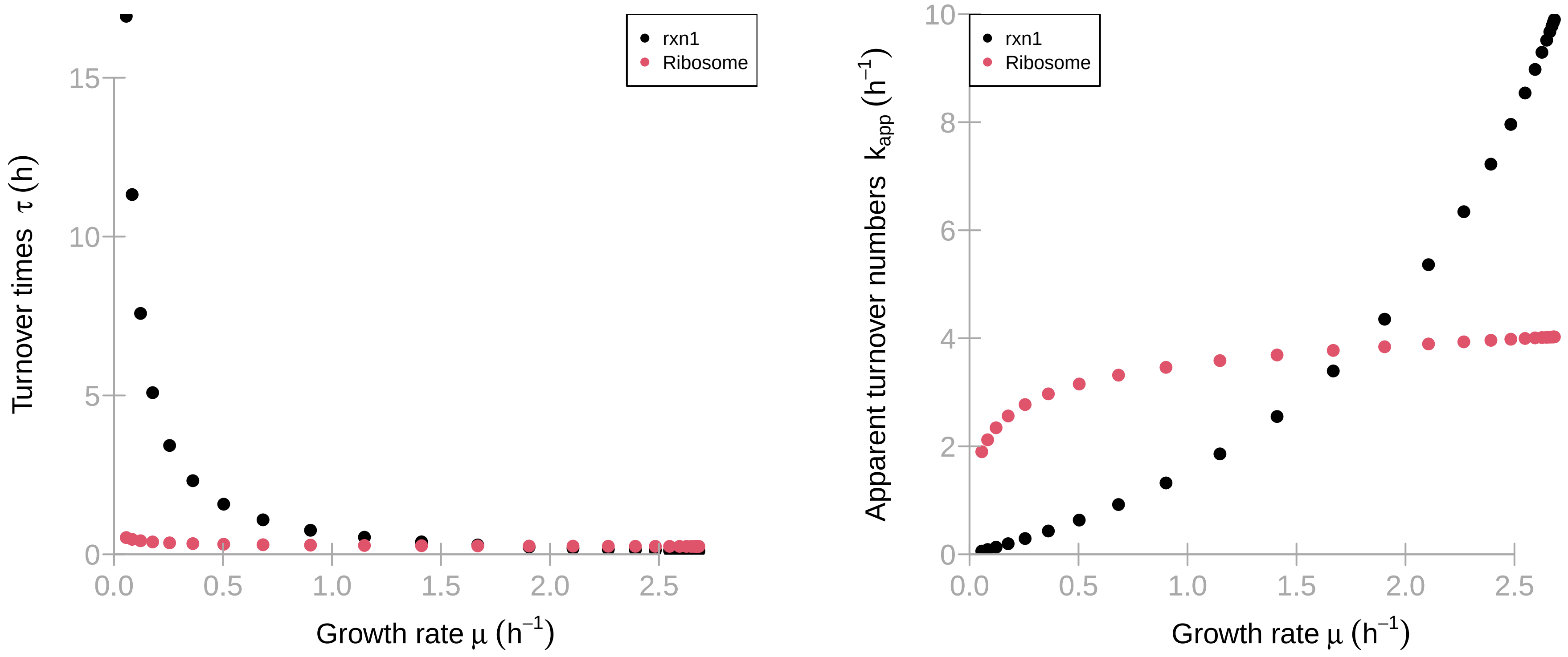

### Model B, mean time (0.0272s) results.pdf

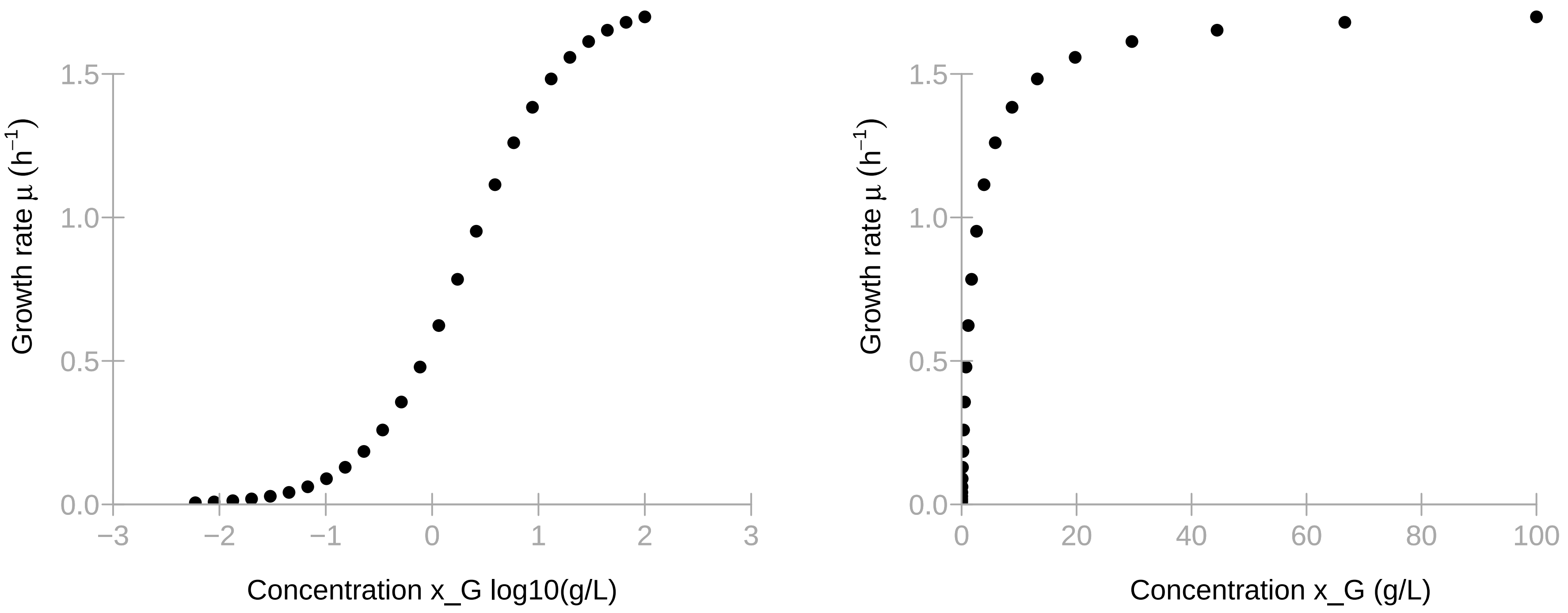

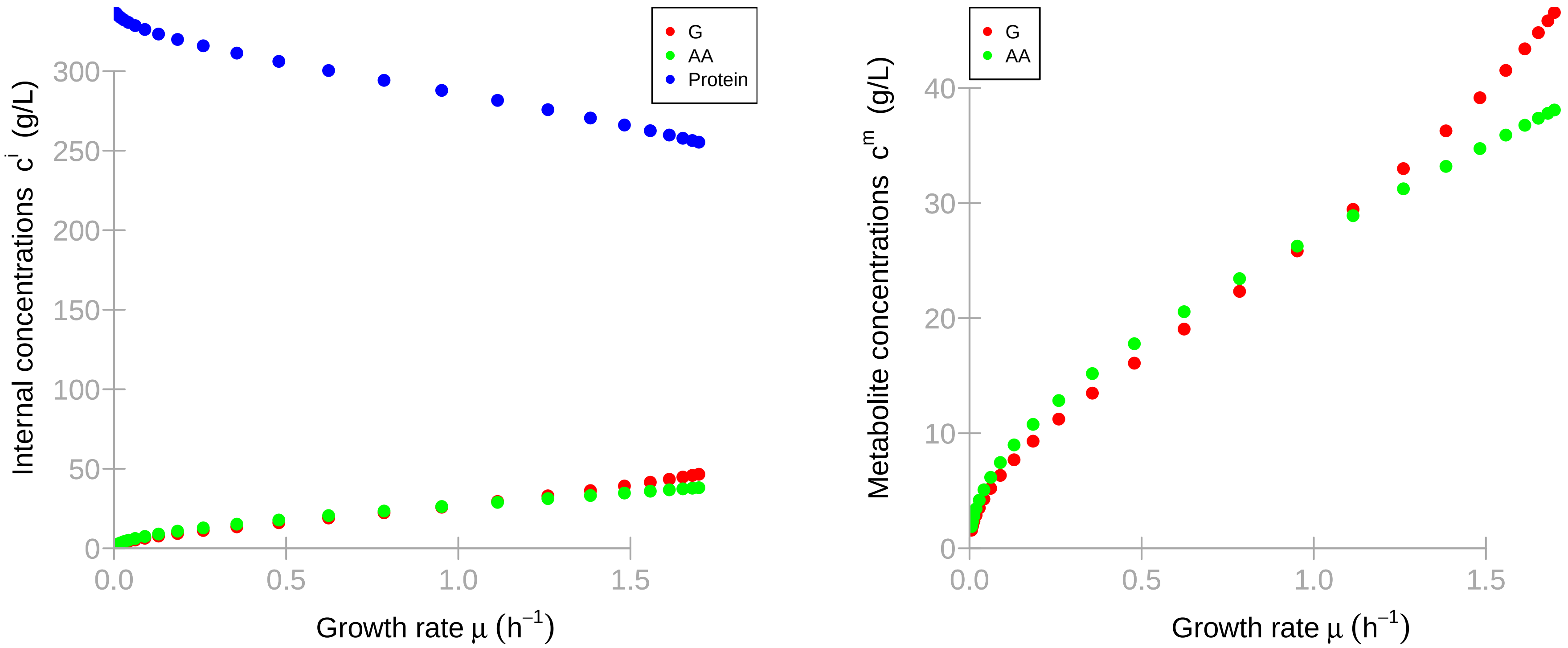

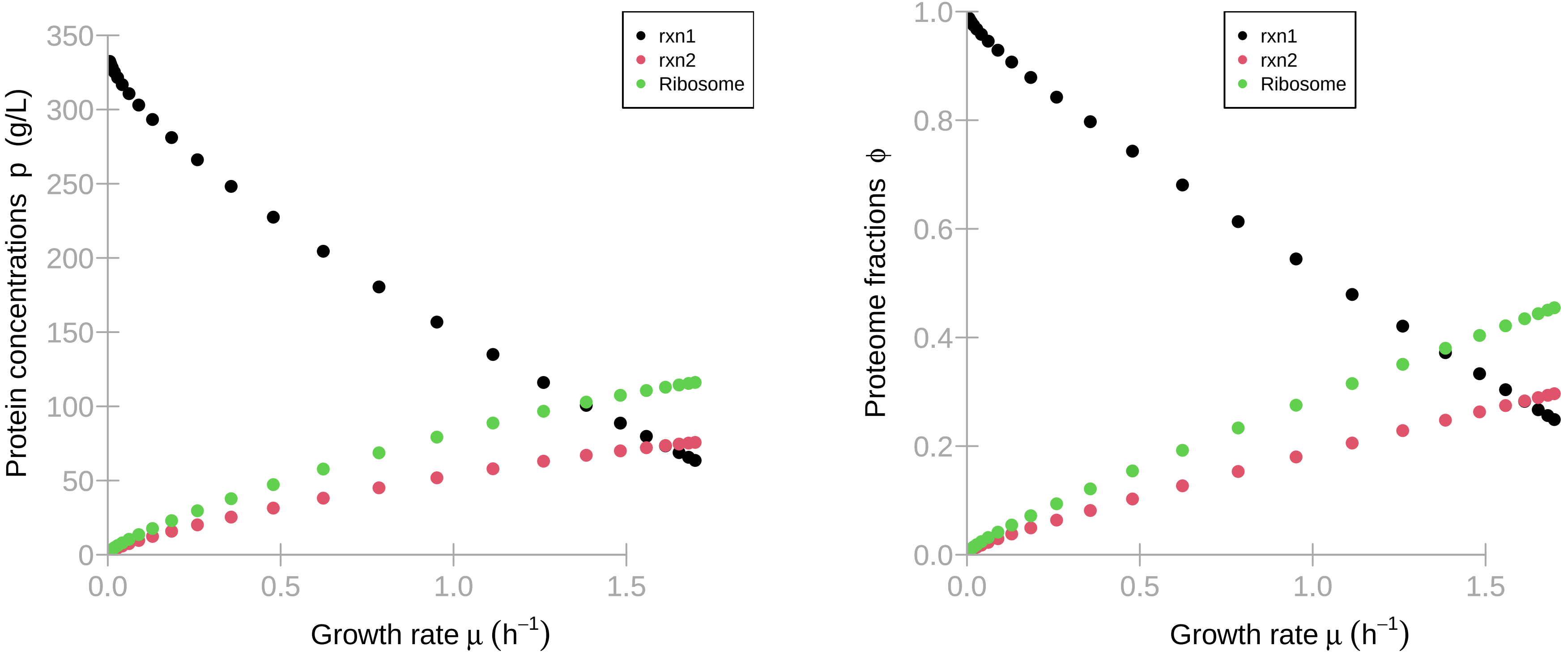

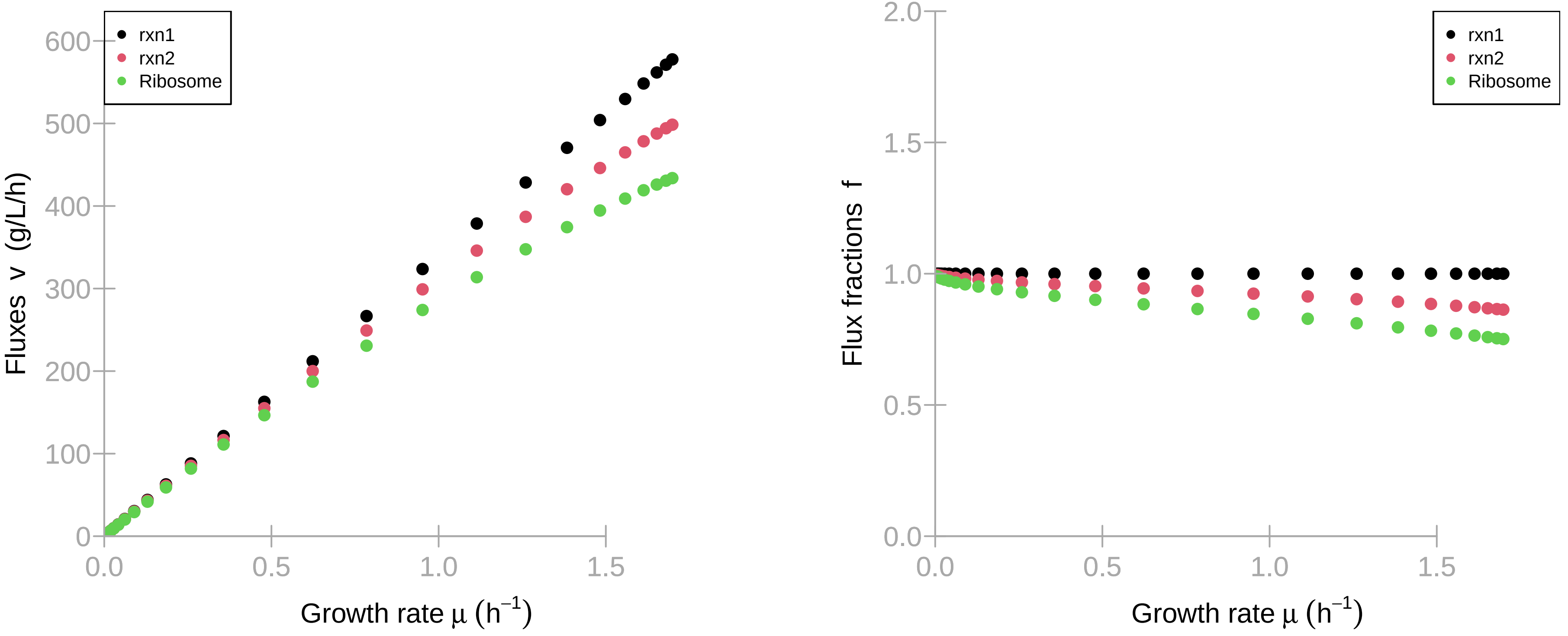

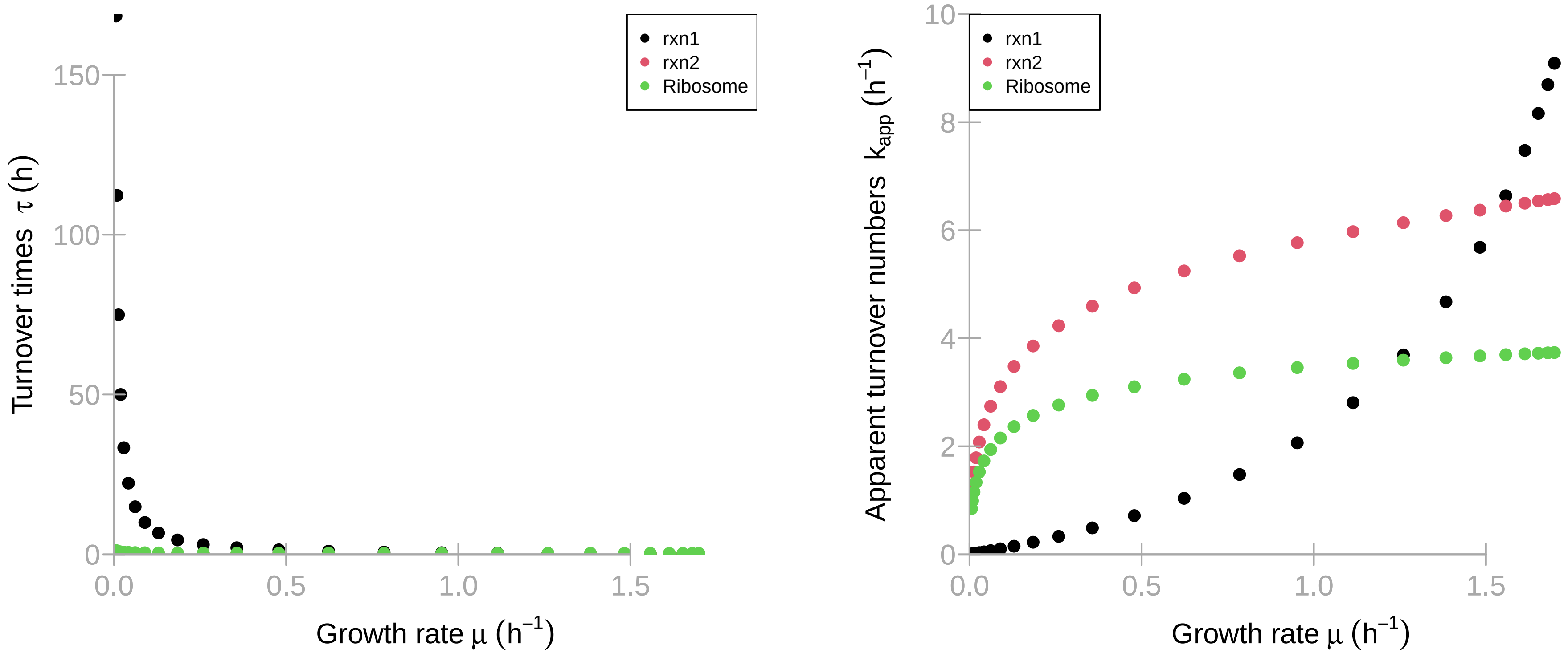

### Model C, mean time (0.0372s) results.pdf

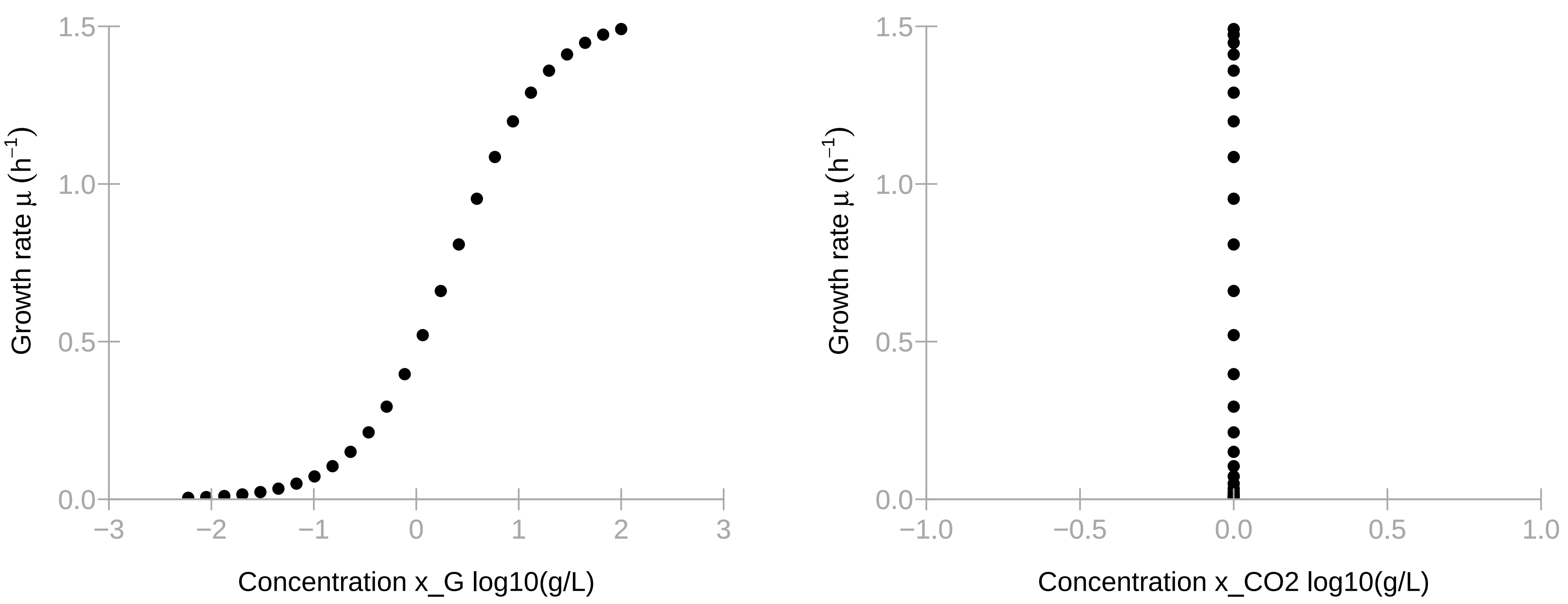

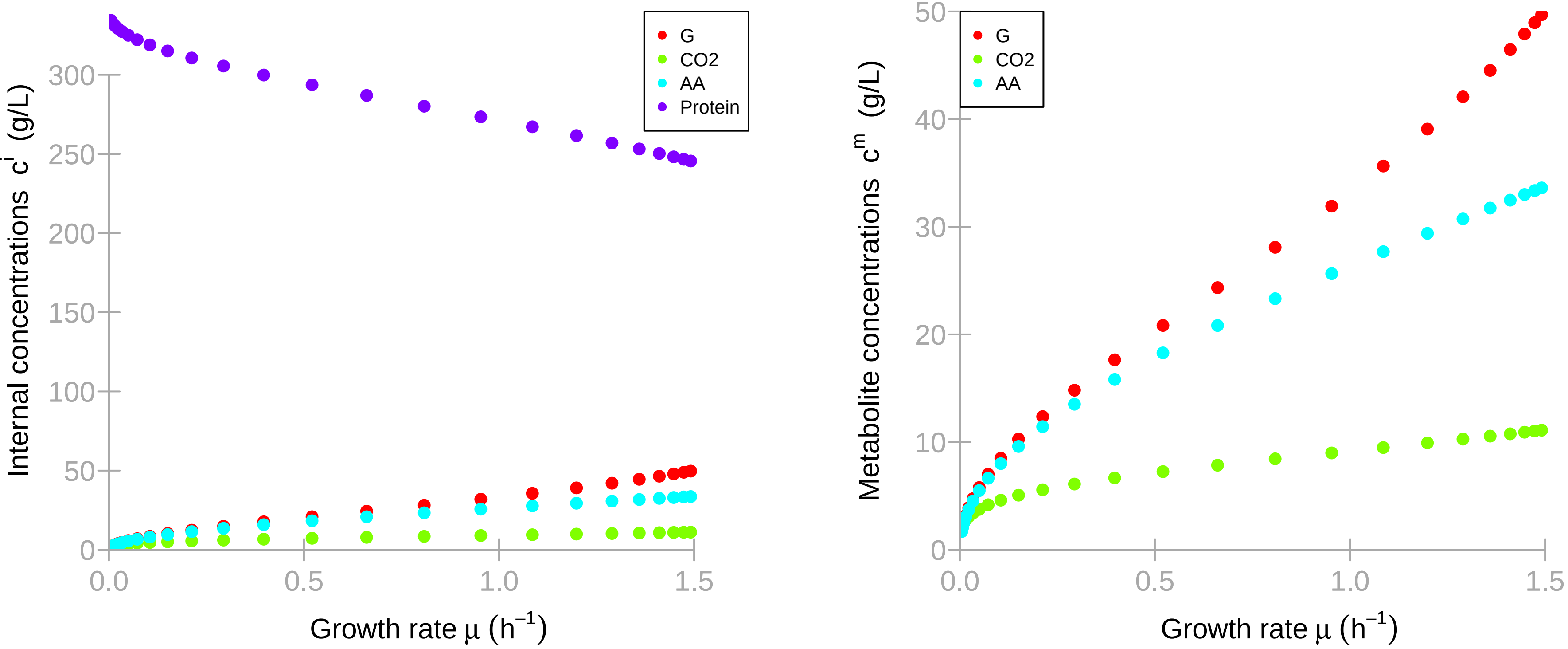

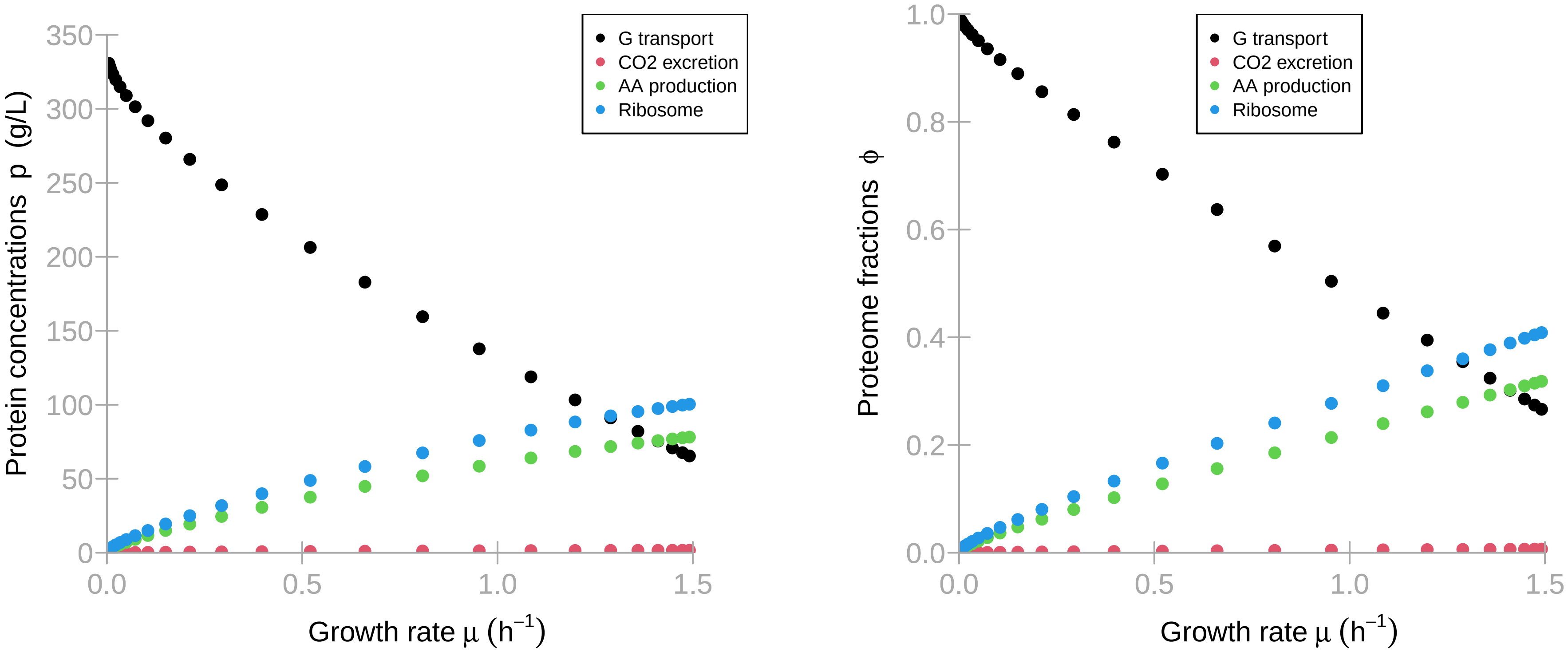

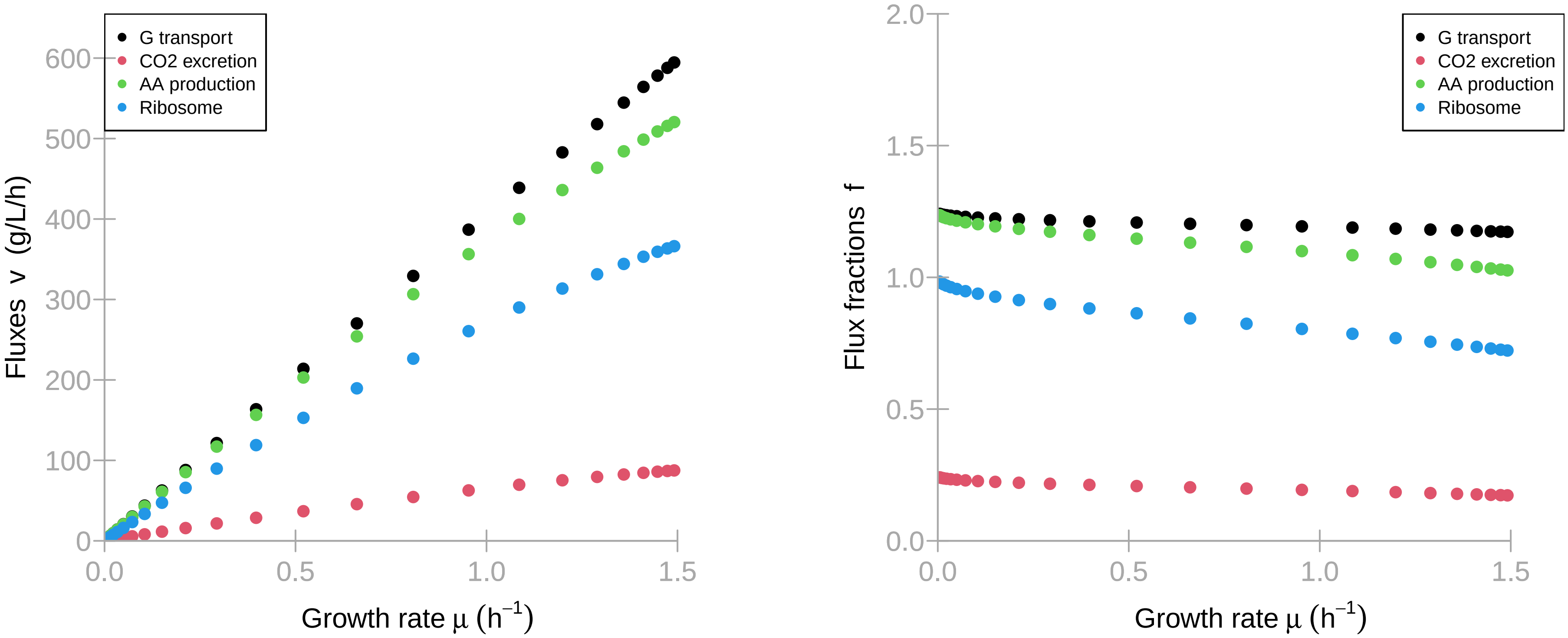

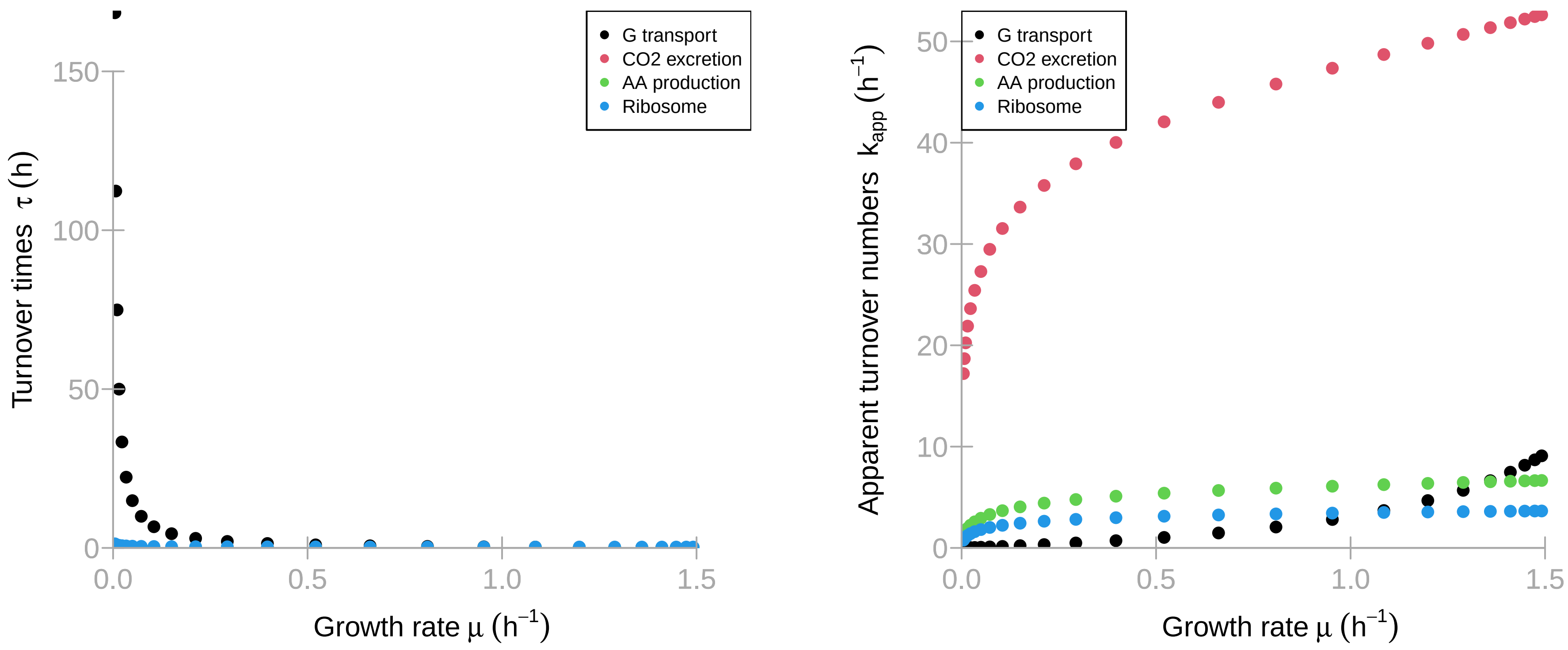

### Model D, mean time (0.08s) results.pdf

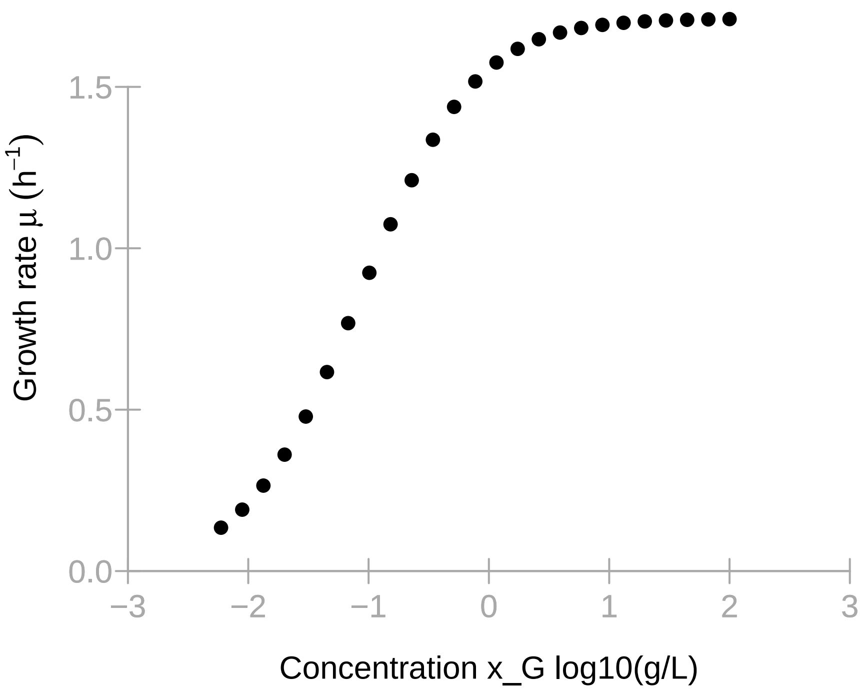

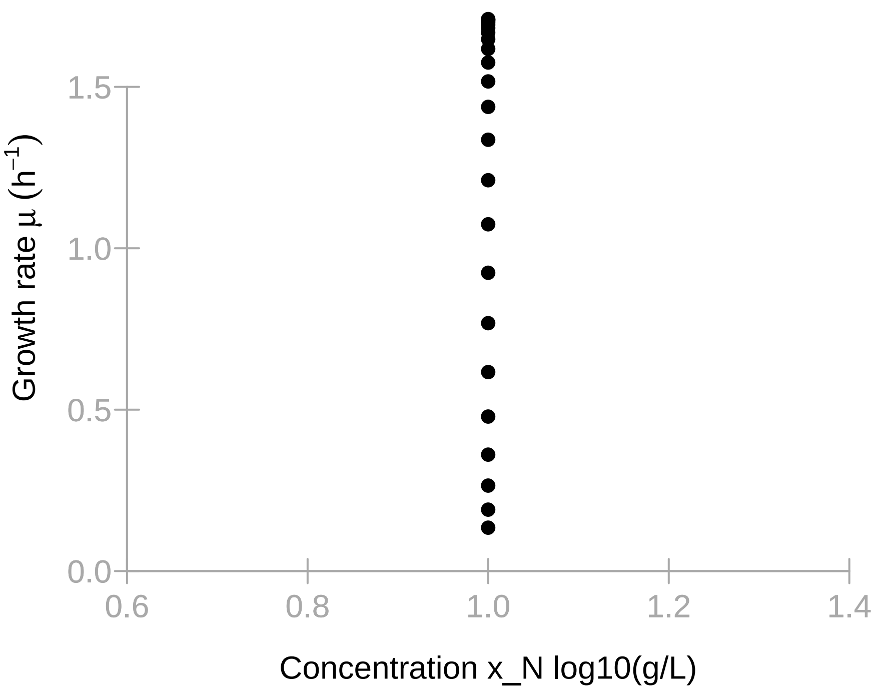

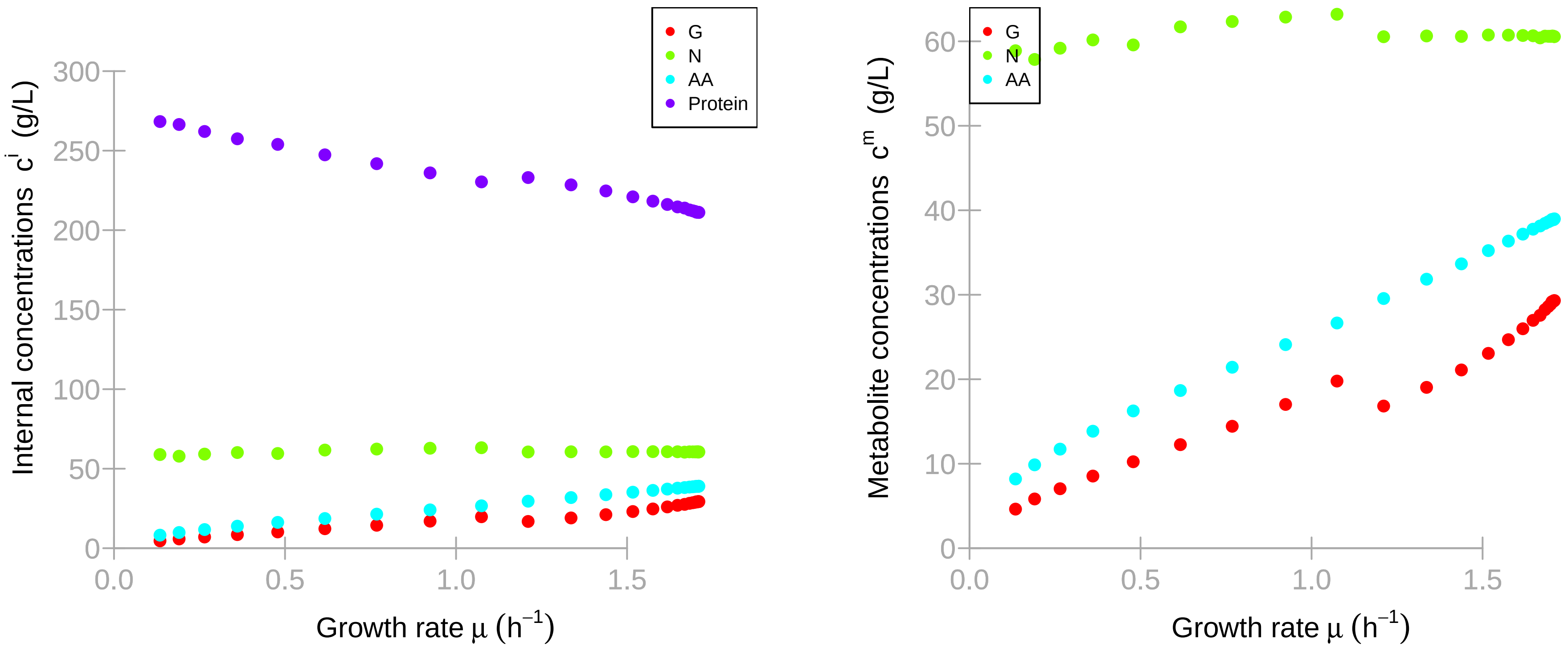

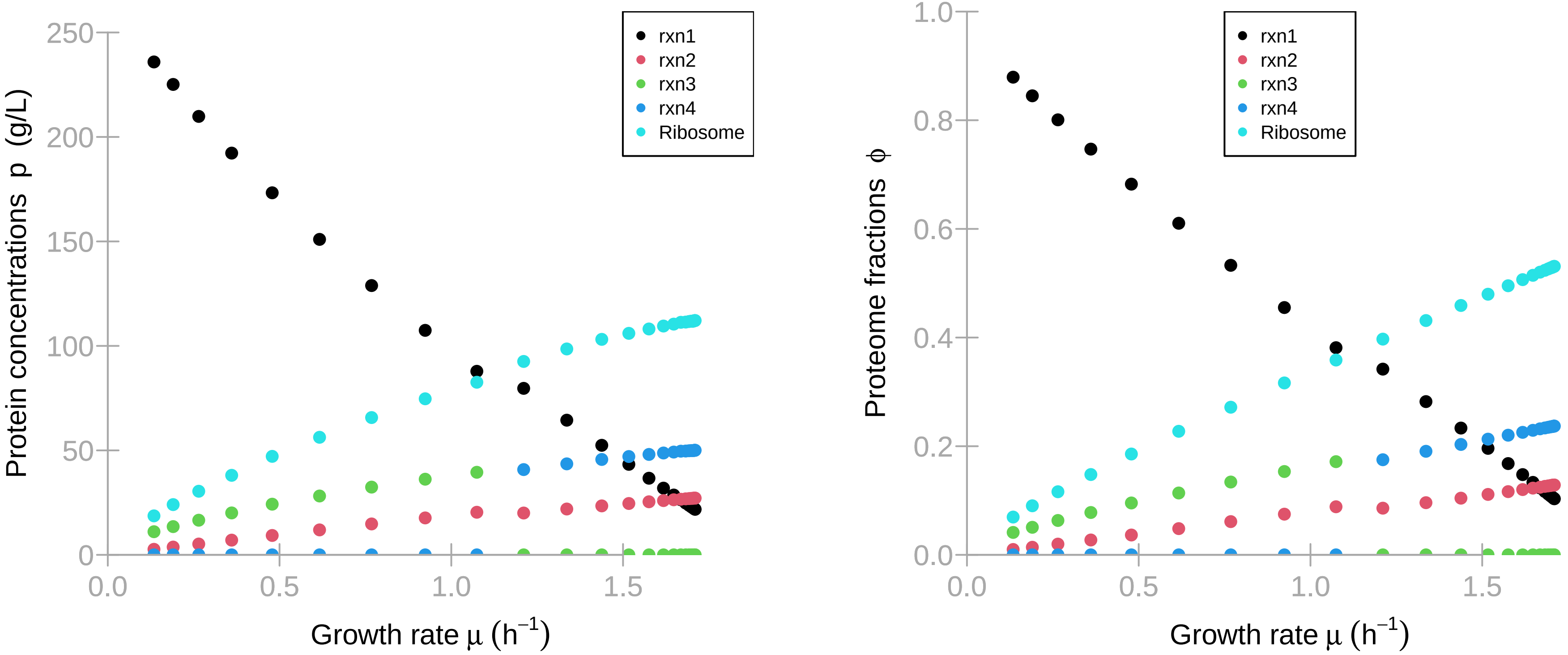

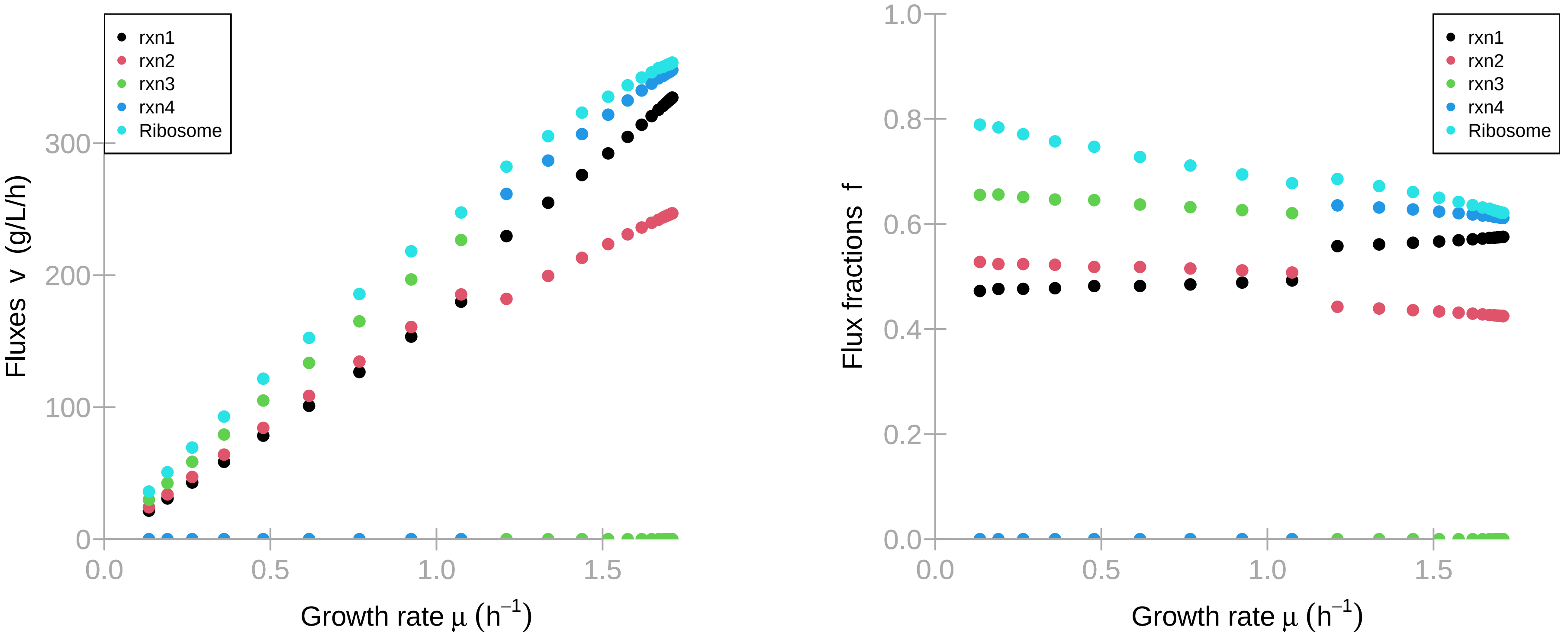

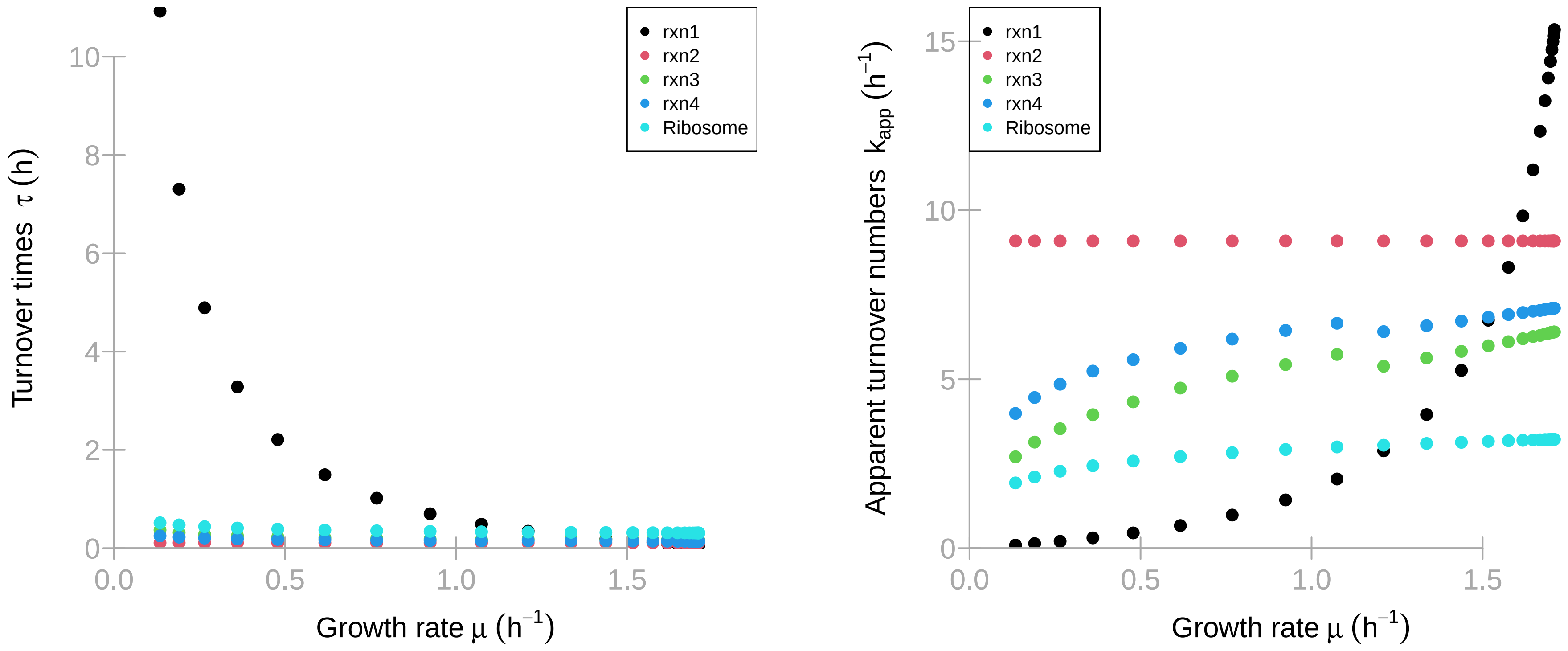
